# Host-microbe interactions in the chemosynthetic *Riftia pachyptila* symbiosis

**DOI:** 10.1101/651323

**Authors:** Tjorven Hinzke, Manuel Kleiner, Corinna Breusing, Horst Felbeck, Robert Häsler, Stefan M. Sievert, Rabea Schlüter, Philip Rosenstiel, Thorsten B. H. Reusch, Thomas Schweder, Stephanie Markert

## Abstract

The deep-sea tubeworm *Riftia pachyptila* lacks a digestive system, but completely relies on bacterial endosymbionts for nutrition. Although the symbiont has been studied in detail on the molecular level, such analyses were unavailable for the animal host, because sequence information was lacking. To identify host-symbiont interaction mechanisms, we therefore sequenced the *Riftia* transcriptome, which enabled comparative metaproteomic analyses of symbiont-containing versus symbiont-free tissues, both under energy-rich and energy-limited conditions. We demonstrate that metabolic interactions include nutrient allocation from symbiont to host by symbiont digestion, and substrate transfer to the symbiont by abundant host proteins. Our analysis further suggests that *Riftia* maintains its symbiont by protecting the bacteria from oxidative damage, while also exerting symbiont population control. Eukaryote-like symbiont proteins might facilitate intracellular symbiont persistence. Energy limitation apparently leads to reduced symbiont biomass and increased symbiont digestion. Our study provides unprecedented insights into host-microbe interactions that shape this highly efficient symbiosis.

## Introduction

All animals are associated with microorganisms (Bang et al., 2018; Bosch and McFall-Ngai, 2011), and consequently, animal-microbe interactions shape life on our planet. While research has concentrated for decades on pathogenic associations, beneficial, i.e. mutualistic symbioses are increasingly moving into the center of attention (McFall-Ngai et al., 2013).

Mutualistic relationships are often based on nutritional benefits for both partners: Symbionts supply their host with nutrients otherwise lacking in the hosts’ diet, while the host in turn provides the symbionts with metabolites, shelter and optimal growth conditions (Moya et al., 2008). To establish and stably maintain their alliance, the partners have to interact on the molecular level. The hosts’ immune system needs to control the symbiont population without erasing it altogether (Feldhaar and Gross, 2009), for example by restricting the symbionts to certain organs and/or by down-regulating its own immune response (reviewed in Nyholm and Graf, 2012). Symbionts, on the other hand, often employ strategies resembling those of pathogens to colonize and persist in their host. For example, similar protein secretion systems are employed by both, symbionts and pathogens, for interactions with the host (Dale and Moran, 2006; Hentschel et al., 2000; McFall-Ngai, 2008; Moya et al., 2008).

In many animals, host-microbe interactions are difficult to assess, due to the high number of potentially involved microbes and the presence of long- and short-term associations, which are hard to distinguish (McFall-Ngai, 2008). Therefore, low-complexity models are important to identify and characterize interaction mechanisms (Webster, 2014). Symbioses of marine invertebrates and their chemoautotrophic symbionts have emerged as suitable study systems. In these symbioses, animal hosts such as gutless annelids and bivalves are often tightly associated with one or few symbiont types, which enable the eukaryotes to prevail in otherwise hostile environments (Dubilier et al., 2008). One of the most conspicuous representatives of these associations, and the first animal in which chemoautotrophic symbionts were discovered, is the giant tube worm *Riftia pachyptila* (short *Riftia*) that thrives around deep-sea hydrothermal vents of the East Pacific (Cavanaugh et al., 1981; Felbeck, 1981). The host’s absolute dependency on its symbiont makes *Riftia* an ideal system to study beneficial host-microbe interactions in a mutualistic symbiosis.

The worm completely lacks a digestive system, but instead receives all necessary nutrients from one phylotype of chemosynthetic endosymbionts (Cavanaugh et al., 1981; Distel et al., 1988; Hand, 1987; Robidart et al., 2008). The host in turn provides the endosymbionts with all necessary inorganic compounds for chemosynthesis (Stewart and Cavanaugh, 2005). This association is remarkably productive: *Riftia* grows extraordinarily fast (> 85 cm increase in tube length per year, Lutz et al., 1994) and reaches body lengths of up to 1.5 m (Jones, 1981).

The uncultured gammaproteobacterial *Riftia* symbiont, tentatively named *Candidatus* Endoriftia persephone (Robidart et al., 2008), densely populates bacteriocytes in the host trophosome, a specialized organ that fills most of the worm’s body cavity (Hand, 1987). The bacteria oxidize inorganic reduced compounds such as hydrogen sulfide to generate energy for carbon fixation (Cavanaugh et al., 1981; Fisher et al., 1989; Markert et al., 2011; Petersen et al., 2011; Robidart et al., 2011; Van Dover, 2000). The symbiont can store elemental sulfur, an intermediate of sulfide oxidation, in sulfur globules (Pflugfelder et al., 2005). Trophosome tissue containing high amounts of stored sulfur has a light yellowish color. During sulfide limitation, i.e., when energy availability is restricted, stored sulfur is consumed and the trophosome appears much darker (Pflugfelder et al., 2005; Wilmot Jr. and Vetter, 1990). The energetic status of the symbiosis can thus be directly inferred from the color of the trophosome.

*Riftia* has been extensively studied, especially with respect to its anatomy, biochemistry, symbiont transmission, and substrate transfer between host, symbionts and the environment (e.g. Drozdov and Galkin, 2012; Liu et al., 2017; Robidart et al., 2011; Sanchez et al., 2007b; Scott et al., 2012; see Stewart and Cavanaugh, 2005 for a review). The symbiont’s metabolism has been studied in detail as well (Stewart and Cavanaugh, 2005), in particular by means of metagenomics and metaproteomics (Gardebrecht et al., 2012; Markert et al., 2007, 2011; Robidart et al., 2008). Yet, little is known about interactions between the two symbiotic partners and, particularly, about the proteins directly involved in these processes.

Our study aimed to illuminate the underlying mechanisms of host-symbiont interactions on the protein level. For this purpose, we employed a state-of-the-art global metaproteomics approach, which required comprehensive sequence data for both partners. While the genome of the *Riftia* symbiont was sequenced previously (Gardebrecht et al., 2012; Robidart et al., 2008), up to now no such information was available for the host. Therefore, we sequenced the transcriptome of the *Riftia* host *de novo*. This enabled us to build a comprehensive protein database, which we used to compare protein abundance patterns in symbiont-containing and symbiont-free *Riftia* tissues. By comparing sulfur-rich and sulfur-depleted specimens, we furthermore examined how host-symbiont interactions vary under high- and low energy conditions. Our analysis sheds light on metabolite exchange processes between both partners, on the host’s symbiont maintenance strategies and on the symbiont’s molecular mechanisms to persist inside the host.

## Results and Discussion

### Interaction analysis of a chemosynthetic deep-sea symbiosis

We sequenced the *Riftia* host transcriptome *de novo* and combined it with three existing symbiont genomes to create a comprehensive holobiont database for identification of *Riftia* host and symbiont proteins (see Material and Methods). Our metaproteomic analysis included comparisons between symbiont-containing and symbiont-free tissues of specimens with light and dark trophosomes (hereafter referred to as sulfur-rich, S-rich specimens and tissues, and S-depleted specimens and tissues, respectively). A fully replicated dataset and stringent experimental design enabled us to find statistically significant differences in individual protein abundance between sample types, as well as abundance differences between functional protein groups. For an overview of all identified proteins, see Supplementary Results and Discussion Part 1 (SOM1). We identified symbiosis-specific proteins and molecular interaction processes, including (i) metabolite exchange between host and symbiont, (ii) host strategies of symbiont maintenance, and (iii) symbiont mechanisms to persist inside the host. Furthermore, we found that (iv) S availability affects symbiotic interactions in *Riftia*. For a graphical representation of the main interactions, see Figure 1. Beyond the results presented here, our data sets also provide a valuable resource for future *Riftia* studies and microbe-eukaryote symbiosis research in general.

**Figure 1:**
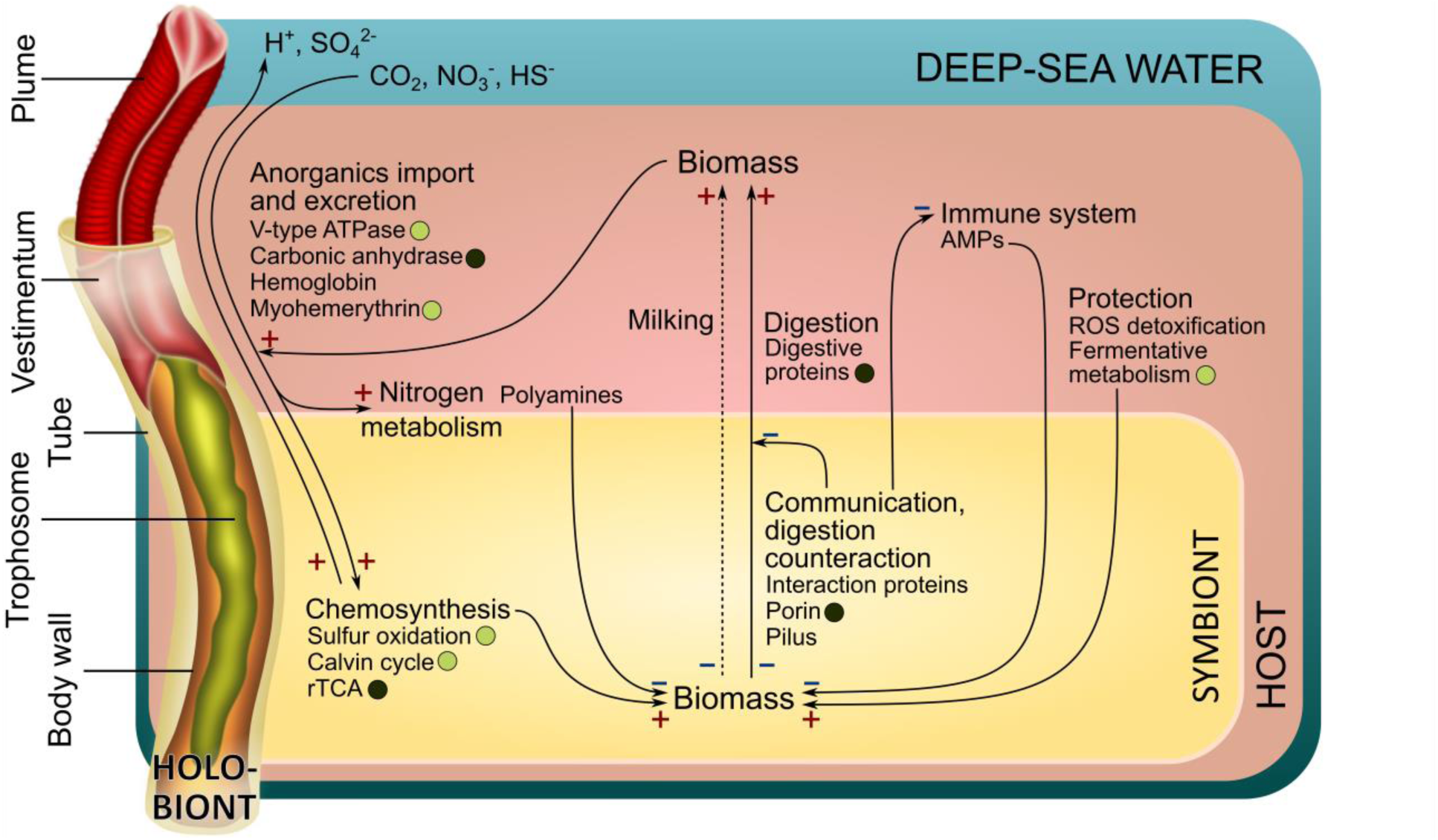
Main interactions in the *Riftia* symbiosis. “+” indicates presumably stimulating interactions, “-“ indicates presumably inhibiting interactions. Circles, where present, indicate that the respective proteins are more abundant in S-rich (light circles) or S-depleted (dark circles) specimens, respectively. Milking: Transfer of small organic compounds (see SOM3).

### Metabolite exchange between host and symbiont

#### *Riftia* digests its symbionts for nutrition

Our results suggest that the main mode of nutrient transfer from symbiont to host is the active digestion of symbiont cells, and that this process might involve endosome-like maturation of symbiont-containing vesicles. We detected a total of 113 host enzymes involved in protein-, amino acid- and glycan degradation as well as in glycolysis and fatty acid beta oxidation. 22 of these proteins were significantly more abundant in trophosome samples than in the other tissues (Table 1). Overall, nearly all of the respective protein groups had higher abundances (i.e. summed-up %orgNSAF) in the symbiont-bearing trophosome than in other tissues, both in S-rich and S-depleted specimens (Figure 2). Many of the protein degradation-related proteins contain signal peptides and are thus likely either contained in lysosomes or secreted into the symbiont-containing vesicles to digest the symbiont cells (Table 1, Supp. Table S1).

**Figure 2:**
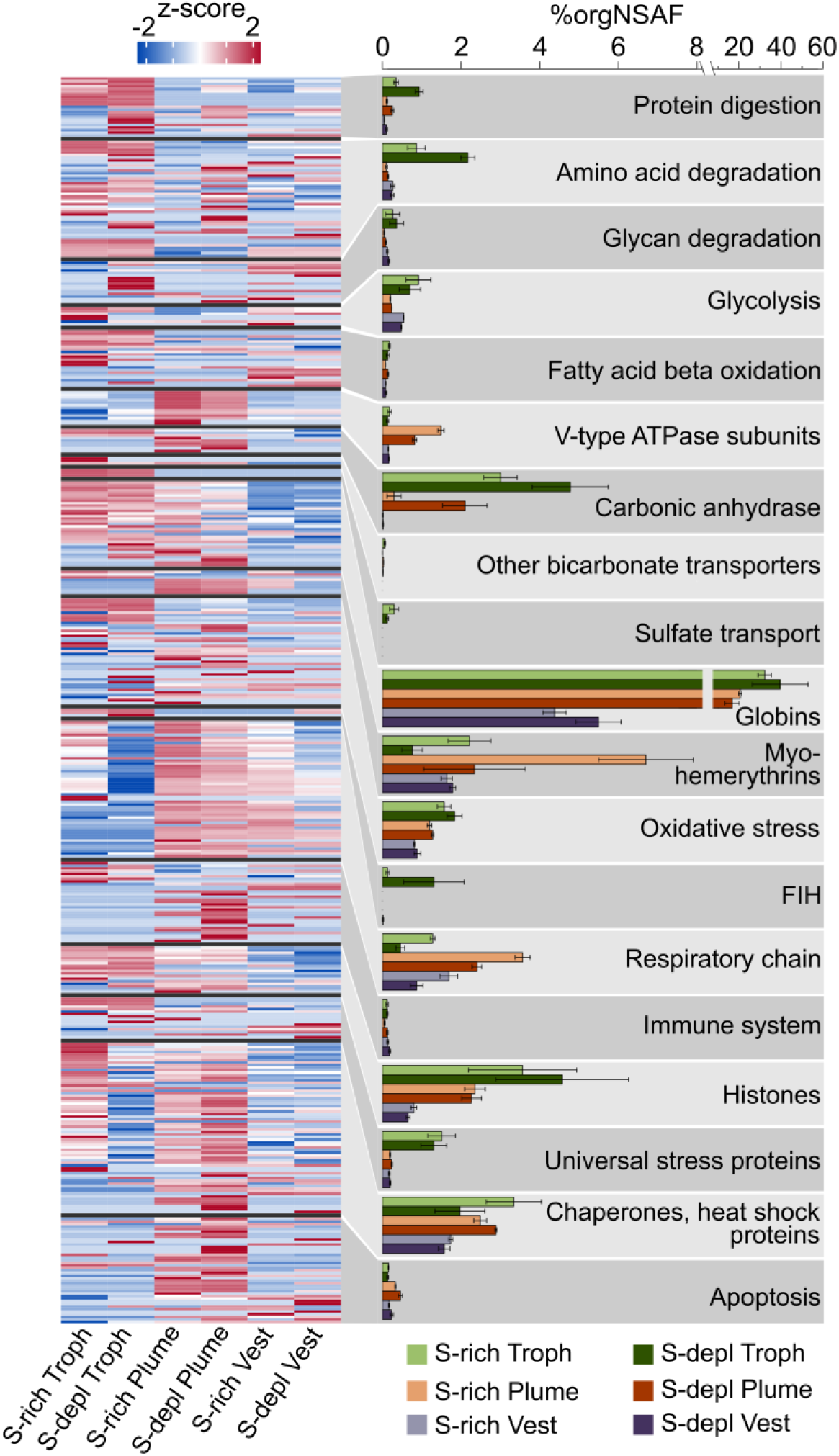
Functional groups of selected *Riftia* host proteins and their relative abundances in tissue samples. The heatmap shows log-normalized, centered and scaled protein abundances. The bar chart shows summed up abundances (%orgNSAF) of the proteins in the respective category. Error bars indicate standard error of the mean. Note the different scaling in the right part of the x-axis. The “Chaperones, heat shock proteins” category also includes chaperonins and Clp proteases. FIH: factor inhibiting hypoxia-inducible factor 1α. S-depl: S-depleted. To view the list of all identified proteins, including their abundances, see Supp. Table S1. (The table can be filtered for the same main or sub categories as presented in this figure; these categories are labelled with “X” in column “Figure 2”). Vest: vestimentum. Troph: trophosome.

**Table 1:**
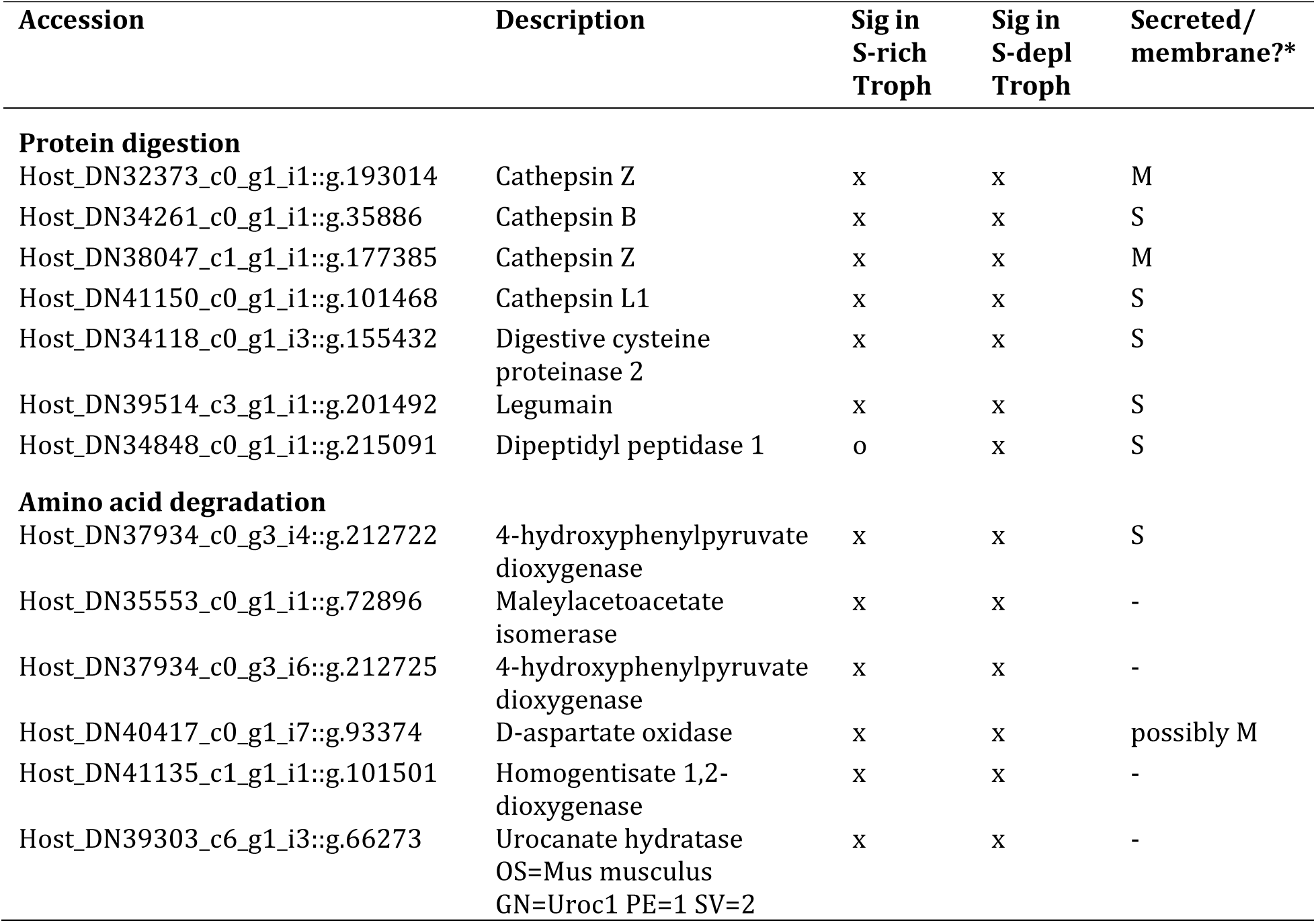
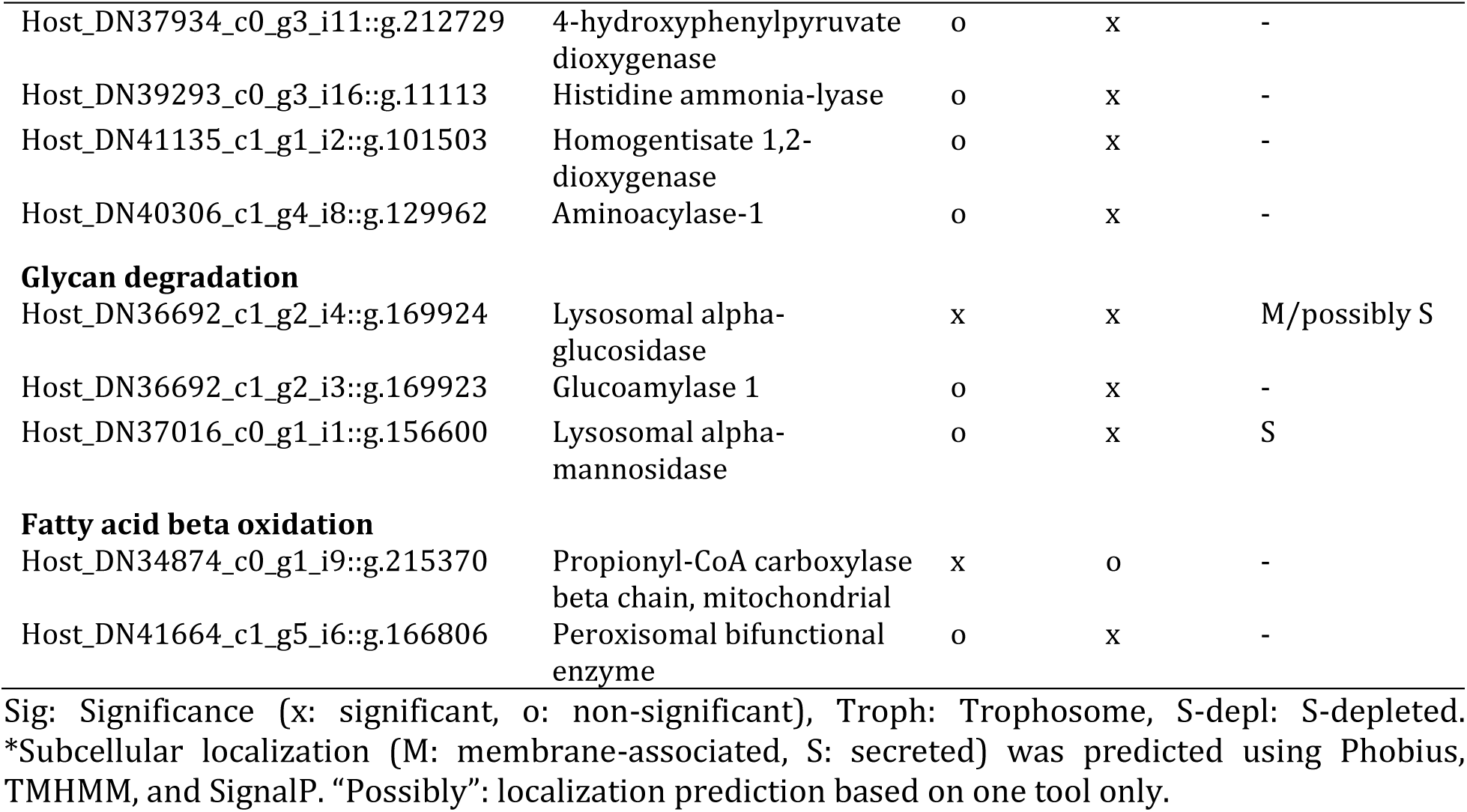
Proteins which are putatively involved in symbiont digestion and which had significantly higher abundances in trophosome samples than in other tissues of S-rich and S-depleted specimens.

Our findings are in accordance with previous biochemical, autoradiographic and microscopic studies, which suggested symbiont digestion in *Riftia* trophosome (Boetius and Felbeck, 1995; Bright et al., 2000; Hand, 1987; Pflugfelder et al., 2009). Moreover, abundant degradative enzymes and symbiont digestion appear to be common in other mutualistic symbioses as well, including deep-sea mussels (Ponnudurai et al., 2017; Streams et al., 1997, Ponnudurai et al., submitted), shallow-water clams (Caro et al., 2009; König et al., 2015) and the gutless oligochaete *Olavius algarvensis* (Wippler et al., 2016; Woyke et al., 2006).

Our metaproteome analysis suggests that symbiont digestion in *Riftia* might involve maturation of symbiont-containing host vesicles in a process resembling the maturation of endosomes. Endosomes form after endocytosis of extracellular compounds and mature from early to late endosomes, which ultimately fuse with lysosomes. The endosome-associated proteins Rab5 and Rab7 showed significantly higher abundances in trophosome samples compared to other host tissues. Rab5 and Rab7 localize to early and late endosomes as well as autophagosomes, respectively, and are markers for these recycling-related organelles (Chavrier et al., 1990; Hyttinen et al., 2013; Vieira et al., 2002). The idea of symbiont degradation via an endosome-like maturation process in *Riftia* is additionally supported by the observation of multilamellar bodies in *Riftia* bacteriocytes in our TEM images (Figure 3). These multilamellar bodies can form in endosomes (Marchetti et al., 2004), but were also suggested to be associated with autophagic cell death in *Riftia* trophosome (Pflugfelder 2009). Although autophagy and apoptosis were suggested to be involved in cell death in *Riftia* trophosome (Pflugfelder et al., 2009), our results contradict this hypothesis. We detected only two autophagy-related proteins (Supp. Table S2) and only 12 of 41 detected apoptosis-related *Riftia* proteins were identified in the trophosome, mostly with similar or significantly lower abundances as compared to other tissues. Caspases, the main apoptotic effectors, were not detected at all on the protein level in trophosome samples (see also SOM2). This is in line with previous microscopic results, which did not indicate apoptosis in the trophosome (Bright and Sorgo, 2003). A non-autophagic, non-apoptotic cell death mechanism was recently described in pea aphid bacteriocytes (Simonet et al., 2018). In the aphids, the proposed mechanism involved hypervacuolation of host bacteriocytes, which was, however, not observed in *Riftia* trophosome. Another caspase-independent cell death mechanism, which involves the protease cathepsin B, has been described in cancer cells (Bröker et al., 2004). As cathepsin B was significantly more abundant in trophosome than in other *Riftia* tissues, we speculate that this protease, amongst other degradative enzymes, may be involved in controlled cell death in *Riftia* trophosome.

**Figure 3:**
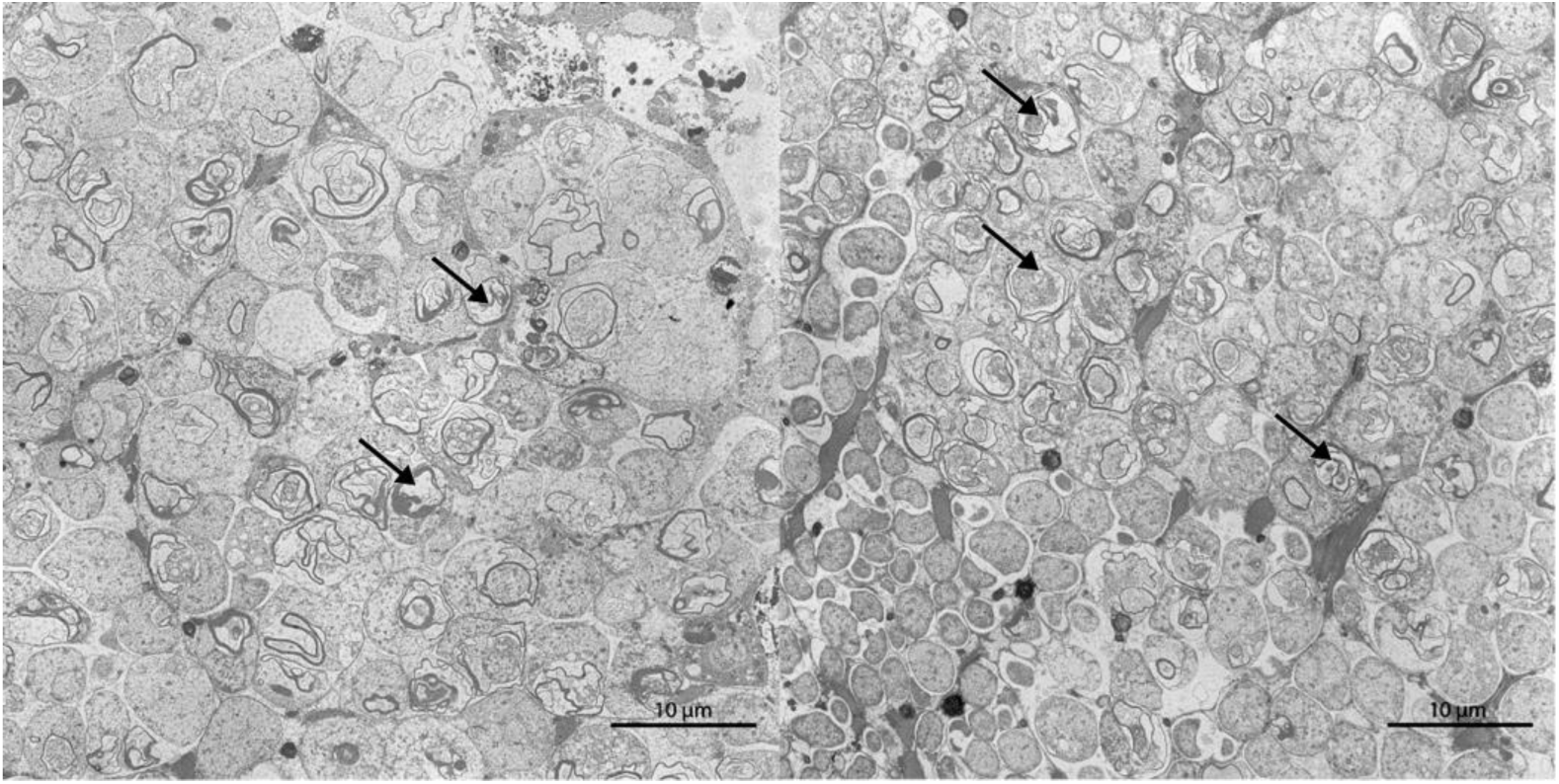
Transmission electron micrographs of *Riftia* trophosome tissue sections. Cell degradation is indicated by the presence of lamellar bodies (black arrows). Brightness and contrast of the micrographs were adjusted for visual clarity. Scale bar: 10 µm

Besides symbiont digestion, a second mode of nutrient transfer, the release of small organic carbon compounds by intact symbionts, was suggested to be present in *Riftia* (Bright et al., 2000; Felbeck and Jarchow, 1998). Our calculated δ^13^C ratios might support this theory (SOM3).

#### *Riftia* dedicates a substantial part of its proteome to provisioning the symbionts with O_2_, sulfide and CO_2_

We found highly abundant and diverse globins, myohemerythrins, V-type ATPase subunits and carbonic anhydrases in the host proteome (Figure 2), indicating that *Riftia* dedicates a substantial part of its proteome to provisioning the symbiont with all necessary substrates for chemosynthesis.

Globins made up about one third of all trophosomal host proteins and one fifth of the total plume proteome (Figure 2), with extracellular hemoglobins being particularly abundant (in sum 32-40%orgNSAF in trophosome and 17-21% in plume samples). *Riftia* has three distinct extracellular hemoglobins composed of globin chains and, in the case of the hexagonal bilayer hemoglobin, globin linker chains (Flores et al., 2005; Zal et al., 1996, 1998). We detected several of these subunits, including isoforms that are (to our knowledge) hitherto undescribed (Supp. Table S1). *Riftia’s* extracellular hemoglobins have been shown to bind both O_2_ and sulfide (Flores et al., 2005, reviewed in Bailly and Vinogradov, 2005; Hourdez and Weber, 2005). Abundant hemoglobins in the highly vascularized plume therefore ensure efficient uptake of these compounds for transport to the symbionts. The symbionts are microaerophilic (Fisher et al., 1989), and simultaneous reversible O_2_- and sulfide-binding to abundant hemoglobins in the trophosome therefore not only provides the bacteria with chemosynthetic substrates and prevents spontaneous sulfide oxidation, but also protects the symbionts from oxygen. (See SOM4 for hemoglobins as a means of protecting the host from sulfide toxicity and for other sulfur metabolic pathways in the host.) In addition to extracellular hemoglobins, we identified four low-abundance (0.002-0.084%orgNSAF) globins that are probably intracellular and might store O_2_ (SOM5).

Besides hemoglobins, myohemerythrins were detected in all tissues, with particularly high abundances of 6.7%orgNSAF in S-rich plumes. With their comparatively high oxygen-binding capacity (Mangum, 1992), hemerythrins could facilitate oxygen uptake from the environment into the plume, and are possibly also involved in O_2_ storage and intracellular transport in *Riftia*. Moreover, the abundance distribution of the nine detected myohemerythrins suggests a tissue-specific function (SOM6).

V-type ATPase subunits were found with highest total abundances of up to 1.5%orgNSAF in *Riftia* plumes (Figure 2), and almost all of the detected subunits were significantly more abundant or exclusively detected in the plumes. V-type ATPases have a pivotal function in regulating internal pH and CO_2_ uptake (De Cian et al., 2003a) and thus in symbiont provisioning. The high energy demand of V-type ATPase-dependent pH regulation could be met via a relatively higher respiration activity in the plume, as indicated by comparatively high total abundances of respiratory chain proteins (Figure 2), ATP synthase and mitochondrial ribosomes in this tissue. Additionally, carbonic anhydrase (CA), another important enzyme for CO_2_ uptake, was detected in all tissues. While we observed tissue-specific abundance patterns of individual CAs (Supp. Figure S4, SOM7), overall CA abundance was highest in the trophosome (Figure 2). CA facilitates CO_2_ diffusion into the plume by converting it to HCO_3_^-^ (De Cian et al., 2003a; Goffredi et al., 1999), and likely back-converts the HCO_3_^-^ to CO_2_ for fixation by the symbionts in the trophosome. Our analysis suggests that three of the *Riftia* CAs could be membrane-bound (SOM7), which might facilitate CO_2_ diffusion into the bacteriocytes by converting HCO_3_^-^ to CO_2_ in the direct cell vicinity (De Cian et al., 2003b; Sanchez et al., 2007a). Transport of HCO_3_^-^ to the bacteriocytes could be mediated by bicarbonate exchangers, which we identified in trophosome and plume samples.

While carbon for fixation by the *Riftia* symbiont is likely mainly transported in the form of CO_2_/HCO_3_^-^, the host may additionally pre-fix CO_2_ into organic C_4_ compounds which are then transported to the symbiont (Felbeck, 1985). We did identify host phosphoenolpyruvate carboxykinase and pyruvate carboxylase, which could be involved in this process (SOM8).

#### *Riftia’s* nitrogen metabolism depends less on the symbiont than previously assumed

*Riftia* symbionts supply their host not only with carbon and energy sources, but also with ammonium produced by bacterial nitrate reduction (Figure 4, SOM9). However, with regard to the subsequent metabolization of organic nitrogen, the host might be more self-sufficient than previously thought: Previous biochemical analyses suggested that only the symbiont, but not the host, can *de novo* synthesize pyrimidines (Minic et al., 2001) and produce polyamines (Minic and Hervé, 2003). In contrast to those studies, we found the multifunctional CAD protein (carbamoyl-phosphate synthetase 2, aspartate transcarbamoylase, and dihydroorotase), in the *Riftia* host metatranscriptome, suggesting that the host can catalyze the first steps of pyrimidine synthesis. As we did not detect CAD protein on the protein level, expression levels and associated activities in the host are likely rather low, and most of the pyrimidine demand could be satisfied by digesting symbionts. In addition, we found key genes involved in polyamine synthesis in the hosts’ metatranscriptome and partially also detected the respective proteins in the hosts’ metaproteome (Figure 4). Our results suggest that while both *Riftia* symbiosis partners can synthesize spermidine, in fact only the host is able to generate spermine. Host spermidine synthase and spermine synthase were exclusively detected in trophosome samples in our study, suggesting that the polyamines produced by these proteins could have a role in symbiont-host interactions. They could, for example, be involved in restricting the symbiont to its cell compartment, i.e. the bacteriocyte vesicle, as suggested for bacterial pathogens (SOM10). In addition, only the host seems to possess a full urea cycle and might degrade not only its own, but also nitrogen-containing metabolites of the symbiont (SOM9). These results show that the symbiont provides the host with necessary metabolic energy and building blocks for biosynthesis, but that the host has also retained key biosynthetic capacities for N-containing organic compounds.

**Figure 4:**
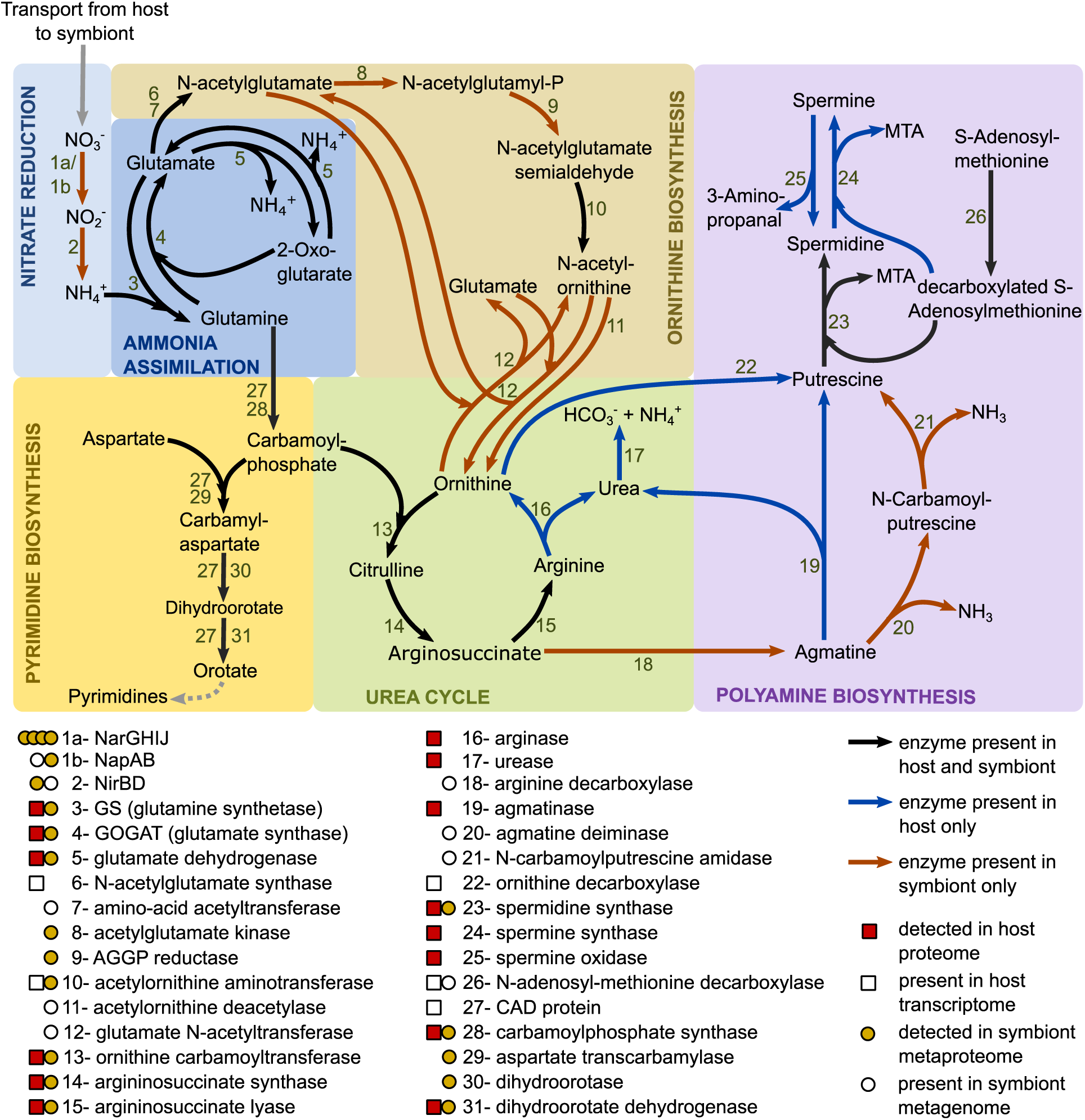
Main nitrogen metabolic pathways in the *Riftia* symbiosis. AGGP reductase: N-acetyl-gamma-glutamyl-phosphate reductase, CAD protein: multifunctional carbamoyl-phosphate synthetase 2, aspartate transcarbamoylase, and dihydroorotase protein, MTA: 5’-methylthioadenosine. Note that the symbiont might also be capable of nitrate respiration (Hentschel and Felbeck, 1993; Markert et al., 2011), which is not depicted here.

### Host strategies of symbiont maintenance

#### *Riftia* protects its symbiont from oxidative damage and may even generate hypoxic conditions in the trophosome

We found several reactive oxygen species (ROS)-scavenging enzymes (superoxide dismutase, peroxiredoxin, glutathione S-transferase), as well as proteins indicative of anaerobic metabolism and universal stress proteins with significantly higher individual abundance and in higher total amounts (summed %orgNSAF) in the trophosome compared to other tissues (Figure 2, SOM11). *Riftia*’s ROS-detoxifying enzymes probably not only protect the host, but also the microaerophilic symbiont against ROS. Upregulation of host proteins involved in ROS detoxification was previously shown in the *Wolbachia* symbiosis (Brennan et al., 2008; Zug and Hammerstein, 2015). Additionally, malate dehydrogenase was highly abundant in trophosomes. This enzyme is regularly observed in different invertebrates under anaerobic conditions (Hourdez and Lallier, 2007) and is involved in maintaining redox balance during anaerobiosis (Fields and Quinn, 1981). The trophosome might thus rely more on fermentative metabolism than on respiration, as also indicated by the overall lower abundance of host respiratory chain proteins in trophosome compared to other tissues of both, S-rich and S-depleted specimens. We also detected hypoxia-inducible factor 1-alpha inhibitors (factor inhibiting HIF1a; FIH) almost exclusively in trophosome samples, which further supports the idea that free oxygen concentrations in the trophosome are low. This is in line with the high oxygen-binding capacity of *Riftia* hemoglobins (Fisher et al., 1989; Hentschel and Felbeck, 1993), and with the suggestion of fermentative metabolism under hypoxic and even oxic conditions in *Riftia*, based on biochemical results (Arndt et al., 1998). Taken together, lower oxygen concentration in the trophosome, (partial) anaerobic host metabolism, and host ROS-detoxifying enzymes in this tissue would not only protect the symbionts from oxidative damage, but would additionally decrease the competition between the *Riftia* host and its symbionts for oxygen.

#### The *Riftia* immune system might be involved in symbiont population control

We detected several proteins which are potentially involved in a specific immune reaction of *Riftia* against its symbiont in the trophosome. Two bactericidal permeability-increasing proteins (BPIPs) were detected, one exclusively in the trophosome, the other only in the plume. BPIPs act specifically against Gram-negative bacteria, causing initial growth arrest and subsequent killing due to inner membrane damage (Elsbach and Weiss, 1998). In *Riftia*, BPIPs could be involved in keeping the symbiont population under control, e.g. as part of the digestion process or by preventing the symbionts from leaving their intracellular host vesicles. Likewise, in the *Vibrio*-squid symbiosis, BPIPs have been implied in restricting the symbiont population to the light organ (Chen et al., 2017). In addition to BPIPs, a pathogen-related protein (PRP) was present in all replicates of S-rich trophosome, but absent from all other tissues. In plants, pathogen-related proteins accumulate during defense responses against pathogens (reviewed in Van Loon and Van Strien, 1999). Pathogen-related proteins have also been described in nematodes (Asojo et al., 2005) and humans (Eberle et al., 2002), although their function remains elusive.

We also found that histones had overall higher abundance in *Riftia* trophosome than in other tissues. Four of these histones were significantly more abundant in trophosomes than in other tissues, and three additional histones were exclusively detected in trophosome samples (Supp. Table S1). Besides being crucial for DNA interactions, histones and histone-derived peptides can have antimicrobial effects (Cho et al., 2009; Park et al., 1998; Rose et al., 1998). A blastp search of the detected *Riftia* histones against the Antimicrobial Peptide (AMP) Database APD3 (Wang et al., 2016) gave hits for four of the *Riftia* histones (Supp. Table S3), stimulating the speculation that these histones may have antimicrobial properties. While AMP-like histone-derived peptides in the plume might be involved in defense against environmental microbes, the high abundance of histones in the trophosome could point to a function in host-symbiont interaction. Host-derived AMPs could, for example, be involved in controlling the symbiont’s cell cycle. In their life cycle, the symbionts apparently differentiate from actively dividing stem cells into growing, but non-dividing larger cells (Bright and Sorgo, 2003). As various AMPs were shown to inhibit cell division or septum formation and to cause filamentous cell morphologies (reviewed in Brogden, 2005), we speculate that *Riftia* AMPs may inhibit cell division as well, e.g. via interaction with symbiont GroEL. Interaction between a host AMP and a symbiont GroEL has been proposed to lead to cell elongation of bacterial weevil symbionts (Login et al., 2011). A role of histones and histone-derived peptides in immune system responses has been described or suggested in various other organisms, including catfish (Park et al., 1998), Komodo dragons (Bishop et al., 2017), toads (Cho et al., 2009) and humans (Rose et al., 1998).

Beyond individual immune system proteins, we did not observe a general immune response of *Riftia* against its symbiont (which is not surprising, as the symbionts are contained inside host vesicles). This indicates that the host immune system does not play a major role in controlling symbiont population size. More likely, symbiont population control might to a large part be a result of digestion of symbionts (a “mowing” process), which effectively prevents the symbionts from escaping their compartments and/or overgrowing the host. Nevertheless, the immune system might be involved in phage protection and symbiont recognition during establishment of the symbiosis (SOM12).

### Symbiont persistence mechanisms

#### Eukaryote-like protein structures in the symbiont might be involved in host communication

The metagenome of the *Riftia* symbiont *Ca.* E. persephone encodes several protein groups with possible roles in symbiont-host interactions, including eukaryote-like protein (ELP) structures, as revealed by our SMART analysis (Supp. Table S4). We detected more than 100 of these symbiont proteins in the trophosome samples (Figure 5), which points to a symbiosis-relevant function.

**Figure 5:**
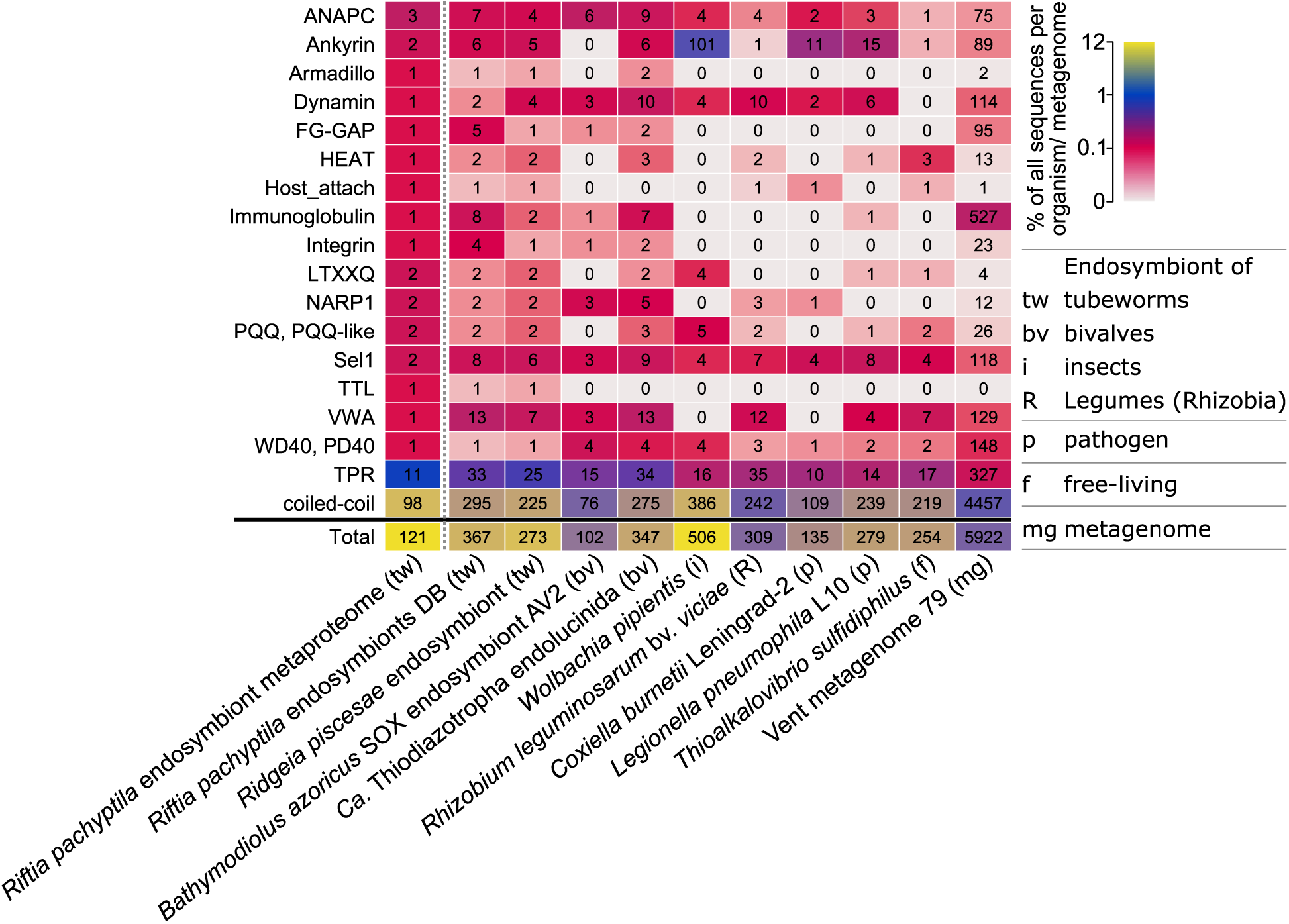
Selected domains with eukaryote-like structures and with putative functions in symbiont-host interactions in the *Riftia* symbiont and in selected other organisms and metagenomes. Color scale: percentage of genes/proteins containing the respective domain relative to all gene/protein sequences in this organism or metagenome. Numbers: total number of genes/proteins containing the respective domain. For an overview of all analyzed organisms and domains see Supp. Figure S5. For details on the organisms see Supp. Table S5. For further information about the selected protein groups see Supp. Table S4. ‘*Riftia pachyptila* endosymbiont metaproteome’ refers to the *Riftia* symbiont proteins detected in this study.

Among the ELPs detected in the symbiont metaproteome were two ankyrin repeat-containing proteins, which contain a signal peptide and are therefore likely secreted (predicted by Phobius, http://phobius.sbc.su.se/). Ankyrin repeats were found to mediate protein-protein interactions (Li et al., 2006). In the sponge *Cymbastela concentrica*, symbiont ankyrins were proposed to interact with the eukaryote’s phagocytosis system: The symbiont ankyrins were heterologously expressed in *E. coli* and led to inhibition of phagocytosis by amoebae (Nguyen et al., 2014). Likewise, a secreted *Legionella pneumophila* ankyrin protein apparently interferes with host endosome maturation (Pan et al., 2008). The *Ca.* E. persephone ankyrin repeat-containing proteins could therefore directly interact with host proteins as well, e.g. to modulate endosome maturation, and thus to interfere with symbiont digestion by the host. Similarly, proteins with tetratricopeptide repeat (TPR)/Sel1 domains, which we also detected in the *Ca.* E. persephone metaproteome, have been shown to impact phagocytosis by amoeba (Reynolds and Thomas, 2016).

The *Riftia* symbiont furthermore encodes eukaryote-like proteins of the tubulin-tyrosine ligase family (TTL proteins). These proteins post-translationally modify tubulin and thus interact with the eukaryotic cytoskeleton (Prota et al., 2013). We found one TTL protein in the *Ca*. E. persephone metaproteome. Other protein groups which are involved in protein-protein interactions in eukaryotes, e.g. with cytoskeletal proteins, and which we detected in *Ca.* E. persephone, include armadillo repeat proteins (Coates, 2003) and HEAT repeat-containing proteins (Yoshimura and Hirano, 2016). As several of the protein structures analyzed here are also found in other mutualistic symbionts and pathogens (SOM13, Supp. Table S4), it is conceivable that parallels exist between interaction processes of mutualistic and pathogenic associations, and that the *Riftia* symbiont employs a strategy similar to that of pathogens to communicate with its host on the molecular level.

#### Symbiont membrane proteins may export effector proteins into host cells and lead to strain adaptation

We detected various outer membrane-related proteins in the *Ca.* E. persephone proteome, including a porin (Sym_EGV52132.1), which was one of the most abundantly expressed symbiont proteins, and 12 type IV pilus (T4P) system proteins (PilQ, PilF, PilC, PilBTU, PilM, PilN, PilP, FimV, PilH, PilY1). Five additional T4P structure proteins were encoded in the metagenome (*pilVWXE*, *pilO*). These proteins are in direct contact with the host cells, and therefore likely involved in interactions between both symbiosis partners, including such processes that facilitate the symbiont’s persistence inside the host cells.

The abundant symbiont porins could transport effector molecules, e.g. to modulate digestion by the host. A role of porins in effector transport during symbiosis has been hypothesized for the *Vibrio fischeri* OmpU, a channel protein that is important for symbiont recognition by the squid host (Nyholm et al., 2009).

The T4P system is a complex structure, which, in *Pseudomonas aeruginosa*, comprises more than 40 proteins, including structural and regulatory proteins (Leighton et al., 2015). It can have several functions in different species: adhesion, secretion and natural transformation (Davidson et al., 2014; Hager et al., 2006; Leighton et al., 2015; Stone and Kwaik, 1999). As the *Ca.* E. persephone T4P system is likely not involved in adhesion to host cells during symbiosis (although it might be during the initial infection), it could participate in protein secretion and/or natural transformation. The *Riftia* symbiont’s T4P system could export putative effector proteins (e.g. ankyrins, SET domain proteins, SOM13, SOM14) for host interactions. Interestingly, in the pathogen *Francisella tularensis* ssp. *novicida*, a T4P structure is involved in secretion of infection-moderating proteins (Hager et al., 2006).

Besides their putative function in effector protein export, symbiont membrane proteins may also lead to bacterial strain adaptation. The *Riftia* symbiont population is polyclonal, i.e. consists of several distinct strains (Polzin et al., 2019). T4P system-mediated exchange of genetic material between different symbiont strains would add to this diversity in the symbiosis and might additionally enable exchange of symbiosis-related genes within the free-living *Ca.* E. persephone population. Natural transformation in symbionts has only recently been shown for *V. fischeri* in culture (Pollack-Berti et al., 2010) and the earthworm symbiont *Verminephrobacter eiseniae*, which likely employs a T4P structure for DNA uptake (Davidson et al., 2014). As microbial cell densities are comparatively high in eukaryote-prokaryote mutualisms, natural transformation in these systems might actually be more common than previously recognized. The proposed DNA uptake by the *Riftia* symbiont may not only facilitate exchange between symbiont strains, but may also promote horizontal gene transfer between host and symbiont, e.g. of eukaryote-like proteins. This hypothesis, as well as the speculation that *Ca.* E. persephone might be capable of conjugation (SOM14) certainly warrant further investigations.

### S availability affects symbiotic interactions in *Riftia*

#### S-depleted *Riftia* hosts digest more symbionts than S-rich specimens

We compared the metaproteomes of *Riftia* specimens with and without stored sulfur (i.e., energy-rich vs. energy-depleted specimens) to examine how energy availability impacts symbiotic interactions. Metabolite transfer is apparently especially influenced by the energy regime: The host supposedly relies more on symbiont digestion in times of S shortage. Proteinaceous symbiont biomass was notably lower in S-depleted trophosomes (32%) than in S-rich trophosomes (58%; Figure 6). Simultaneously, overall abundances for several groups of host digestive enzymes were higher in S-starved trophosomes (Figure 2), and a number of individual host proteins were significantly more abundant in these S-depleted samples, such as enzymes involved in protein digestion (including cathepsin B), amino acid degradation, the late-endosome protein Rab7 and histones (Supp. Table S1). One reason for this supposed increase in symbiont digestion in S-depleted trophosomes could be a lower nutritional value of the energy-depleted symbionts. S-depleted symbionts have lower abundances of enzymes involved in sulfur oxidation, probably due to lower S availability. Therefore, less energy might be available for biosynthesis under S depletion, rendering the symbiont less “nutritious” for the host. S-depleted hosts may thus have less energy available, despite increased symbiont digestion. This idea is supported by the observation that host proteins involved in the energy-generating glycolysis, TCA cycle, respiratory chain, ATP synthesis and biosynthetic pathways were less abundant in S-depleted trophosomes than in S-rich trophosomes. Potentially, increased symbiont digestion under S-depleted conditions is necessary for the host to satisfy its basal metabolic demand. Concomitant with the postulated lower nutritional value of S-depleted symbionts, the Calvin cycle key enzyme RubisCO had an about 10-fold lower abundance in S-depleted symbionts. Abundance of the rTCA cycle key enzyme ATP citrate lyase (EGV51152.1), on the other hand, was slightly higher in S-depleted symbionts than in S-rich symbionts, albeit only 1.4-fold. Under S-depleted conditions, symbionts apparently rely relatively more on the rTCA cycle, which is more energy-efficient than the Calvin cycle (Markert et al., 2007). The Calvin cycle could be used in addition to the rTCA cycle under favorable conditions to maximize overall carbon fixation. Moreover, symbiont enzymes involved in translation were overall more abundant in S-rich trophosomes than in S-depleted trophosomes. Less protein biosynthesis in S-depleted symbionts would not only impact the nutritional value of these symbionts, but additionally directly decrease the proteinaceous symbiont biomass. The reason for the lower proteinaceous biomass of symbionts in S-depleted trophosomes is therefore probably two-fold: The host digests more symbionts and the symbionts produce less biomass compared to energy-rich trophosomes.

**Figure 6:**
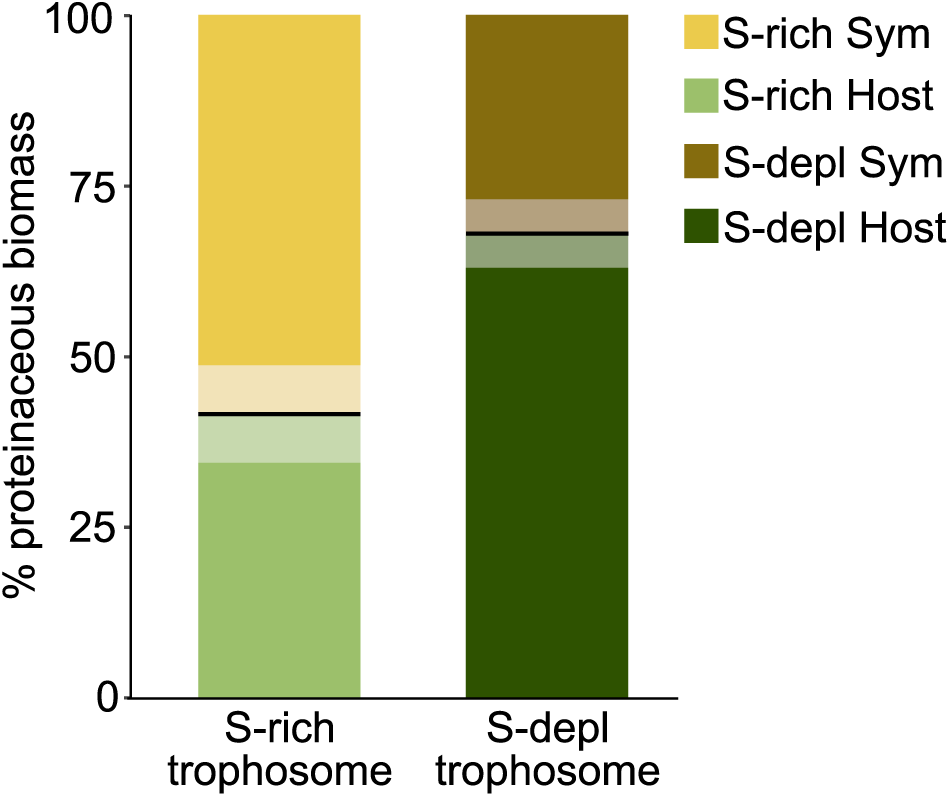
% proteinaceous biomass contributions of host and symbiont as calculated from our metaproteomics results (Kleiner et al., 2017). Bold lines indicate the mean, semitransparent areas indicate standard error of the mean. Sym: Symbiont, S-depl: S-depleted

These findings are in contrast to previous results (Scott et al., 2012), which showed no significant differences in autotrophic activity and symbiont abundance between *Riftia* specimens from high- vs. low-sulfide habitats. Possibly, increased symbiont digestion is a short-term adaptation to fluctuating environmental conditions, whereas under long-term low-S conditions the symbiosis might adapt e.g. by reduced growth rates. Decrease in symbiont abundance or total protein under energy-limiting conditions has also been noted in *Bathymodiolus* (Stewart et al., 2005) and *C. orbicularis* bivalves (Caro et al., 2009) as well as in *O. algarvensis* oligochaetes (Wippler et al., 2016). Relying on the symbionts as nutrient source also under unfavorable conditions thus appears to be a common symbiosis mechanism, which would ensure survival of the host and a subset of the symbiont population, ultimately prolonging survival of the individual holobiont.

#### S availability influences CO_2_ uptake, pH regulation and O_2_ regime in the *Riftia* host

S-depleted hosts seem to invest relatively more biosynthetic capacities in CO_2_ uptake and less in pH regulation, and their trophosomes are supposedly less hypoxic than those of S-rich hosts (SOM11, SOM15). At the same time, S availability appeared to have little influence on non-symbiont-related processes in the host, as only very few (i.e. < 10) individual proteins significantly differed in abundance between S-rich and S-dark plume and vestimentum samples. This indicates that the host’s metabolism is very well buffered against changes in environmental conditions.

#### Higher digestion pressure might result in symbiont countermeasures

In S-depleted *Riftia* specimens, a putative *Ca.* E. persephone dodecin was significantly more abundant than in S-rich specimens. This protein might be involved in protecting the symbiont against oxygen and/or digestion stress (SOM14). A symbiont porin, which was also significantly more abundant in S-depleted specimens, might be involved in counteracting the supposedly higher digestion pressure (see above and SOM14).

## Conclusion

To fully understand the biology of organisms, it is crucial to study them together with their symbiotic partners as holobionts (Gilbert et al., 2012). Given its low complexity, high specificity and extreme dependence of the host on the symbiont, the association of *Riftia* and its bacterial partner serves as an excellent system to study mutualistic host-microbe interactions. While *Riftia* lives in a unique and remote environment, many of the interactions we identified, like symbiont digestion by the host, high host investment in substrate transfer to the symbiont, host-directed symbiont population control, and eukaryote-like symbiont proteins that could interact with the hosts’ molecular machinery, seem to be critical in other symbiotic associations as well, including insects, mussels and oligochaetes. These interactions might therefore represent common principles among evolutionarily diverse mutualistic animal-microbe associations.

Our study provides access to the *Riftia* host transcriptome and protein sequences and thus paves the way for future research on host-microbe interactions in *Riftia* and other systems. Promising research directions include the elucidation of protein functions, e.g. of *Riftia* immune system proteins and symbiont eukaryote-like proteins by heterologous gene expression and biochemical assays in model systems. Moreover, our work stimulates future in-depth studies of the molecular mechanisms involved in recognition of both partners during the initial infection of *Riftia* larvae by free-living symbionts. Putative differences between *Riftia*’s short- and long-term adaptation strategies in response to changing environmental conditions also warrant further investigation.

## Material and Methods

### Sampling

*Riftia* tissue samples were obtained during several research cruises in 2008, 2014 and 2017 with RV Atlantis to the deep-sea hydrothermal vent fields on the East Pacific Rise at 9°50’ N, 104°17’ W. *Riftia* specimens were collected by the human occupied vehicle Alvin or the remotely operated vehicle Jason in approximately 2,500 m water depth. Sampling dates for all *Riftia* tissue samples for proteomics, transcriptomics and transmission electron microscopy (TEM) are summarized in Supp. Table S6. Different specimens were used for proteomics, transcriptomics and TEM. *Riftia* specimens were dissected onboard and tissue samples stored at −80 °C. The lamellae of the tentacular crown were shaved off to provide “plume” samples, trophosome samples were dissected from whole trophosome, body wall samples were retrieved and washed after removal of the trophosome, and vestimental samples were cut off from the lateral portions of the vestimentum. Specimens were classified into sulfur-rich (S-rich), S-depleted and medium S according to their trophosome color (yellow/light green, dark green/black, or medium green, respectively).

### Extraction of whole-tissue RNA

RNA was extracted from a total of 22 tissue samples from 9 specimens (6 x trophosome, 6 x body wall, 5 x plume, 5 x vestimentum, see Figure 1). Tissue samples were homogenized by bead-beating with lysing matrix D (MP Biomedicals) in 1 ml TRIzol® (Thermo Fisher Scientific; 3x 6.5m/s for 30 s, 3 min cooling on ice in between). After 5 min acclimatization to room temperature, samples were applied onto QIAShredder columns (Qiagen) and centrifuged (16,000 x g, 3 min, 4 °C). Afterwards, RNA was isolated from the aqueous flow-through according to the TRIzol extraction protocol, with the modification that samples were centrifuged for 20 min at 12,000 x g and 4 °C for phase separation. Glycogen was added for RNA precipitation. RNA was washed twice with 75% ethanol and purified using the Norgen RNA Clean-Up and Concentration kit according to the manufacturer’s Protocol A, including DNA removal with DNase (Qiagen). Quality of extracted RNA was assessed using Nanodrop (Thermo Fisher Scientific) and Bioanalyzer (Agilent) analyses.

### Transcriptome sequencing and assembly

#### Transcriptome sequencing

Transcriptome sequencing was performed employing the TruSeq stranded mRNA (poly A-based) library protocol (Illumina) on a HiSeq 4000 (Illumina), according to the manufacturer’s guidelines.

#### Transcriptome assembly

High-throughput paired-end Illumina sequencing resulted in an average of about 26 million reads per end per library (min 16,045,121 reads per end, max 31,318,532 reads per end, 95% CI 1,673,590). After de-multiplexing and quality-checking of reads in FastQC v0.11.5 (Andrews, 2010), we trimmed low quality bases and adapters with Trimmomatic v0.32 (Bolger et al., 2014) using the settings ILLUMINACLIP:AllAdapters.fa:2:30:10 SLIDINGWINDOW:4:20, and LEADING:5 TRAILING:5 HEADCROP:15 MINLEN:75. Although bacterial mRNA does not possess a polyA tail, previous research has shown that bacterial reads can still be present in polyA-enriched RNA-Seq libraries (Egas et al., 2012). To filter out potential symbiont contaminations from our host transcriptomes, we used the Bowtie 2 v2.2.9 aligner (Langmead and Salzberg, 2012) in very-sensitive mode to map the quality-filtered paired-end reads against the published genomes of the endosymbionts of *Riftia* (“Riftia1”, NCBI locus tag prefix RIFP1SYM, and “Riftia2”, locus tag prefix RIFP2SYM) and *Tevnia jerichonana* (Gardebrecht et al., 2012). Unmapped paired-end reads were subsequently extracted using SAMtools v1.4.1 (Li et al., 2009). Potential environmental sequence contaminations from sample handling were excluded with DeconSeq v0.4.3 (Schmieder and Edwards, 2011) using coverage and identity thresholds of 0.90 and 0.95, respectively. The decontaminated host reads were normalized and assembled with Trinity v2.3.2 (Grabherr et al., 2011). To optimize the transcriptome assembly we performed four different assemblies with different parameters and input files: 1) only paired reads, 2) paired and unpaired reads, 3) only paired reads plus jaccard-clip option (to reduce chimeras), 4) paired and unpaired reads plus jaccard-clip option.

To assess the completeness of the different assemblies we compared our transcriptomes against the BUSCO v2.0 eukaryote and metazoan orthologous datasets (Simão et al., 2015). Overall, the best results in terms of transcriptome completeness and quality were obtained by the assembly approach using paired and unpaired reads plus jaccard-clip option (Supp. Table S7). This dataset was used for all further analyses.

#### Open reading frame (ORF) prediction

TransDecoder v3.0.1 (Haas et al., 2013) was used to identify coding regions in the assembled transcripts. To improve ORF prediction, we examined all candidate ORFs for homology to known proteins by searching the Swiss-Prot (http://www.uniprot.org) and Pfam (Finn et al., 2016) databases (downloaded January 3, 2017) with BLASTP (Altschul et al., 1990, e-value 1e-05) and HMMER3 (Eddy, 2009), respectively. ORFs that were longer than 100 amino acids and/or had a database entry were retained. The FASTA headers of the TransDecoder output files were modified with a custom PERL script to include the BLASTP protein annotations.

### Database generation

A common database for protein identification of *Riftia* host and symbiont was generated. To this end, host protein sequences were clustered at 95% identity with CD-HIT v. 4.6 (Huang et al., 2010). For symbiont sequences, the three proteomes of the *Riftia*1, *Riftia*2 and *Tevnia* symbiont (Gardebrecht et al., 2012) were used. *Riftia*1 was used as basis for clustering the symbiont protein sequences with CD-Hit-2D (Huang et al., 2010). Subsequently, the combined symbiont database was clustered at 95% identity. Identifier prefixes were added to distinguish between host and symbiont sequences for Calis-p (Kleiner et al., 2018, see below). Host and symbiont databases were concatenated and the cRAP database containing common laboratory contaminants (The Global Proteome Machine Organization) was added. The final database contained 71,194 sequences.

### Proteomics sample preparation and analysis

For metaproteomics analysis, we used three biological replicates per tissue (trophosome, vestimentum, plume) and condition (specimens with S-rich and S-depleted trophosomes), which resulted in a total of 18 samples. Tissues were disrupted by bead-beating for 45 s at 6.0 m/s with lysing matrix D tubes (MP Biomedicals) in SDT buffer (4% (w/v) sodium dodecyl sulfate (SDS), 100 mM Tris-HCl pH 7.6, 0.1 M dithiothreitol (DTT)), followed by heating to 95 °C for 10 min. Tryptic peptides were generated following the FASP protocol of Wiśniewski et al. (2009) with minor modifications as described by Hamann et al. (Hamann et al., 2016). Peptide concentrations were determined with the Pierce Micro BCA assay (Thermo Scientific Pierce) according to the manufacturer’s instructions. The tryptic digest was desalted on-line during LC-MS/MS analysis.

All samples were analyzed by 1D-LC-MS/MS as in Hinzke et al. (2019), using 4 h gradients. Samples were analyzed in a randomized block design (Oberg and Vitek, 2009) and run in technical triplicates. Two technical replicate runs were acquired with a 50 cm analytical column, one with a 75 cm analytical column. To standardize the stable isotope fingerprinting (SIF) analysis (Kleiner et al., 2018), human hair was measured in technical duplicate alongside the *Riftia* samples in the replicate run using a 75 cm column.

### Proteomics data evaluation

#### Protein identification, quantification and statistical analyses

For protein identification, MS/MS spectra of combined technical triplicate runs were searched against the combined host and symbiont database using the Sequest HT node in Proteome Discoverer version 2.0.0.802 (Thermo Fisher Scientific) as in Kleiner et al. (2018). For protein abundance estimates, normalized spectral abundance factors (NSAFs, Zybailov et al., 2006) were calculated per sample and organism (%orgNSAF, Mueller et al., 2010). Statistical evaluation was performed based on spectral counts using the edgeR package (Robinson et al., 2010) in R (R Core Team, 2017). The edgeR package uses an overdispersed Poisson model for analysis of count data. Overdispersion is moderated across proteins using empirical Bayes methods (Robinson et al., 2010). We employed a false-discovery rate (FDR) of 0.05 to assign statistical significance to protein abundance differences. For graphical representation, heatmaps were generated with the R package ComplexHeatmaps (Gu et al., 2016) and intersection plots with the R package UpsetR (Lex et al., 2014). Protein biomasses of host and symbiont were calculated as in Kleiner et al. (2017).

δ^13^C values of *Riftia* symbiont and host were calculated from mass spectrometry data with Calis-p (Kleiner et al., 2018) using one technical replicate LC-MS/MS run (75 cm analytical column). Human hair was used as reference material.

#### Protein annotations, functional characterization and categorization

Besides the annotations included in the database, proteins where further characterized using the online tools described in Supp. Table S8. Proteins were manually categorized into functional groups based on their annotations and the information in the Uniprot (The UniProt Consortium, 2017), NCBI (https://www.ncbi.nlm.nih.gov/) and InterPro (Finn et al., 2017) databases. We used the Transporter Automatic Annotation Pipeline (TransAAP) (http://www.membranetransport.org/transportDB2/TransAAP_login.html) of the TransportDB2 (Elbourne et al., 2017) and TCDB (Saier Jr et al., 2016) with gblast 2 (http://www.tcdb.org/labsoftware.php) to annotate transporters in the *Riftia*1 symbiont metagenome database. To detect possible antimicrobial peptides (AMPs) among the host proteins, we searched the detected host proteins against the antimicrobial peptide database APD3 (Wang et al., 2016) using BLASTP (Altschul et al., 1990) in BLAST+ 2.7.1 (Camacho et al., 2009). Results were filtered for %identity >75% and e-value < 0.005. We screened the *Riftia* proteome for homologs of known autophagy-related *Drosophila melanogaster* proteins (as listed in Chang and Neufeld, 2010) by Blast-searching (BLASTP, Altschul et al., 1990) in BLAST+ 2.8.1, Camacho et al., 2009) the *Riftia* host proteome against the respective *Drosophila* amino acid sequences (Supp. Table S2).

#### SMART analysis of eukaryote-like and potential interaction domains

We used the SMART tool (Letunic and Bork, 2018) to screen the *Riftia* symbiont protein database for proteins and domains which could be involved in symbiont-host interactions. Structures which did not meet the threshold required by SMART were excluded, whereas overlapping features were included. We manually filtered the SMART annotations to find putative interaction-relevant structures based on the Pfam and SMART database information. To compare the *Riftia* symbiont with other host-associated (mutualistic or pathogenic) and free-living organisms, we also included domains not present in the *Riftia* annotations, but possibly relevant for host-bacteria interactions in other organisms based on the literature. All annotations we included are given in Supp. Table S4. The organisms and their proteome accession numbers we used for comparison can be found in Supp. Table S5. Proteins with structures that did not pass the threshold criterion in SMART were removed.

#### Multiple sequence alignments

We used the alignment tool MUSCLE provided by EMBL (https://www.ebi.ac.uk/Tools/msa/muscle/) for multiple sequence alignment of protein sequences. Alignments were verified visually.

### Transmission electron microscopy (TEM)

The trophosome sample for TEM was fixed at room temperature for 1 h in fixative containing 4% paraformaldehyde, 1% glutaraldehyde, 10% sucrose in 50 mM HEPES (glutaraldehyde was added directly before use) and stored at 4 °C. The sample was washed three times with washing buffer (100 mM cacodylate buffer [pH 7.0], 1 mM CaCl_2_, 0.09 M sucrose) for 10 min each step and treated with 1% osmium tetroxide in washing buffer for 1 h at room temperature. After three additional washing steps in washing buffer for 10 min each, the sample was dehydrated in a graded series of ethanol (30%, 50%, 70%, 90%, and 100%) on ice for 30 min each step. Afterwards, the material was subjected to stepwise infiltration with the acrylic resin LR White according to Hammerschmidt et al. (2005). Sections were cut with a diamond knife on an ultramicrotome (Reichert Ultracut, Leica UK Ltd), stained with 4 % aqueous uranyl acetate for 5 min and finally examined with a transmission electron microscope LEO 906 (Carl Zeiss Microscopy GmbH) at an acceleration voltage of 80 kV. The micrographs were edited using Adobe Photoshop CS6.

### Data availability

The mass spectrometry proteomics data and the database have been deposited to the ProteomeXchange Consortium via the PRIDE (Vizcaíno et al., 2016) partner repository with the dataset identifier PXD012439. Transcriptomics raw data have been deposited to the NCBI Sequence Read Archive (https://www.ncbi.nlm.nih.gov/sra) with the BioProject accession number PRJNA534438 (https://www.ncbi.nlm.nih.gov/sra/PRJNA534438). The datasets will be released upon acceptance of the manuscript in a peer-reviewed journal.

## Supporting information

Supplementary Results and Discussion

## Acknowledgements

We thank the captains and crews of RV Atlantis, DSV Alvin, and ROV Jason for their excellent support during the cruises AT15-28, AT26-10, AT26-23, and AT37-12, which were funded through grants of the US National Science Foundation. We are grateful to Ruby Ponnudurai for sampling, to Jana Matulla and Annette Meuche for excellent technical assistance, to Marc Strous for supporting this project by providing access to the proteomics equipment, to Xiaoli Dong for help with database annotations and to Maryam Ataeian, Jackie Zorz and Angela Kouris for help with MS measurements. Sandy Gerschler did preliminary SMART analyses. Målin Tietjen and Lizbeth Sayavedra gave valuable input for RNA sample preparation. This work was supported by the German Research Foundation DFG (grant MA 6346/2-1 to S.M., grant BR 5488/1-1 to C.B.), the German Academic Exchange Service DAAD (T.H.), a fellowship of the Institute of Marine Biotechnology Greifswald (T.H.), the Canada Foundation for Innovation, the Government of Alberta and a Natural Sciences and Engineering Research Council of Canada NSERC through a Banting fellowship (M.K.), the US National Science Foundation (grants OCE-0452333, OCE-1136727, OCE-1131095, and OCE-1559198 to S.M.S), and *The WHOI Investment in Science Fund* (S.M.S). P.R. is supported by a grant from DFG CCGA Comprehensive Center for Genome Analysis, Kiel and the DFG CRC1182 “Origin and function of metaorganisms”. R. H. and T.B.H.R. were supported by the Deutsche Forschungsgemeinschaft (DFG) CRC1182 ‘Origin and Function of Metaorganisms’, Subprojects B2, Z3 & INF.

## Author contributions

T.H., S.M. and M.K. designed experiments, T.H. prepared and analyzed samples for metaproteomics with input from M.K., compiled metaproteomics database with input from M.K. and S.M., performed statistical analyses, prepared samples for RNA sequencing with input from C.B., prepared figures, wrote manuscript. T.S. was involved in project coordination. T.R., R.H. and P.R. coordinated transcriptome sequencing. H.F. helped with sampling. S.M.S. obtained funding for the research cruises and coordinated sampling as chief scientist. C.B. assembled and annotated transcriptomic data. All authors contributed to the final manuscript.

