## Supplementary Results and Discussion for "Host-microbe interactions in the chemosynthetic *Riftia pachyptila* symbiosis"

### 1) General overview

We identified a total of 3,838 *Riftia* host proteins (Supp. Table S1). Many of these proteins showed statistically significant differences in their relative abundance between tissues (see Supp. Table S9 for total numbers of significantly different protein abundances). The “core” proteome of about 700 proteins, which were identified in all host organs under both environmental conditions (i.e. S-rich and S-depleted specimens; Supp. Figure S1), included ribosomal proteins, proteins involved in RNA-, amino acid- and protein metabolism, energy generation, globins, signaling pathways, oxidative stress response as well as cytoskeletal proteins, which are all important throughout the animal. Host proteins only present in the symbiont-bearing trophosome included digestive proteins, stress proteins, sulfate transporters, some histones and proteins involved in amino acid degradation (Supp. Figure S1). Proteins exclusively identified in the plumes included proteins involved in RNA metabolism, apoptosis, carbonic anhydrases, cell division, cytoskeleton and the extracellular matrix, the immune system, signal transduction and mitochondrial ribosomes.

Moreover, we identified a total of 1,016 symbiont proteins in the trophosome samples (Supp. Table S10). Proteins identified in both S-rich and S-depleted symbionts belonged to all main metabolic pathways. Proteins exclusively identified in S-rich symbionts included ribosomal proteins as well as proteins involved in amino acid-, lipid-, and cofactor biosynthesis, nitrogen metabolism, transcription and translation. Proteins only identified in S-depleted symbionts were mostly low-abundant and included mainly hypothetical proteins and a porin. While we did identify a total of 12 symbiont proteins in other tissues than the trophosome, only four of them were present in more than one biological replicate of vestimentum and/or plume samples. These four belonged to the most abundant symbiont proteins (i.e. adenylylsulfate reductase subunits A and B, RubisCO and porin), and are therefore likely contaminations from the dissection procedure. More analyses are necessary to clarify whether the low-abundance proteins are contaminations or proteins secreted by the symbiont.

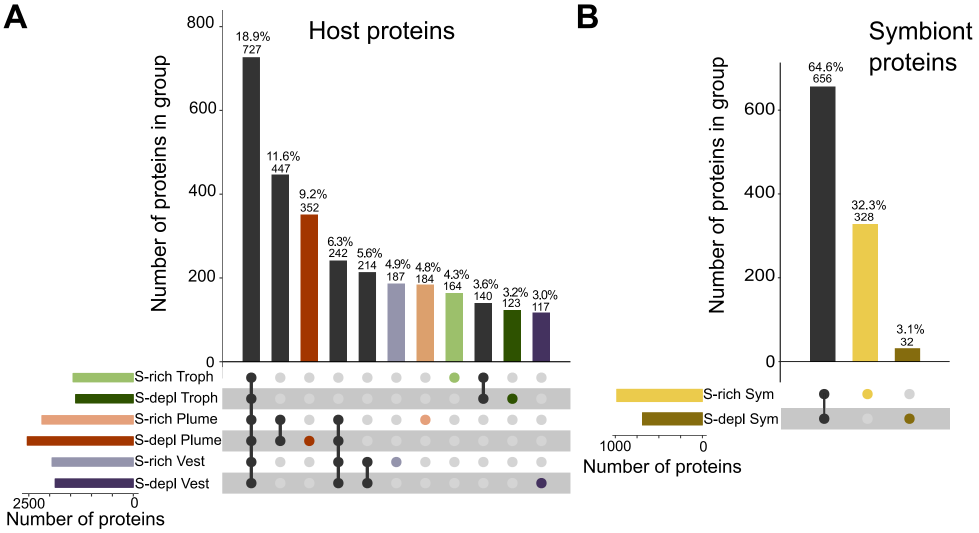


Supp. Figure S1: Overview of the *Riftia* symbiosis metaproteome. A) Intersection plot of the detected *Riftia* host tissue proteins. Bars in the lower left part show the total number of proteins identified in the respective tissue, i.e. in trophosome (Troph), plume or vestimentum (Vest), and condition, i.e. S-depleted (S-depl) or S-rich. Dark/colored dots in the lower right part indicate the samples (single dots) or sample groups (chained dots) in which the proteins are present, equivalent to intersections in a Venn diagram. Columns in the upper part indicate the number and percentage of proteins which are exclusively present in the respective sample or sample group. Shown are selected intersections. B) Intersection plot of *Riftia* symbiont proteins detected in trophosome tissue. Colors in the figures are specific to the respective tissue/symbionts (e.g. yellow for S-rich symbionts, dark green for host proteins from S-depleted trophosome, etc.).

### 2) Stress reactions and apoptosis in *Riftia* tissues

We identified heat shock proteins and other chaperones as well as universal stress proteins (USPs) of the host in comparatively high total abundances in the trophosome. Four USPs were significantly more abundant in trophosome samples than in the other tissues. Heat shock proteins and other chaperones could be involved in protecting host proteins and DNA in the trophosome against reactive oxygen species (ROS) and toxic sulfur compounds. These toxic compounds could be generated in the course of respiration, symbiont sulfur oxidation and/or symbiont digestion by the host. This would be in accordance with observations in *B. azoricus*, where higher HSP70 levels in the gills than the mantle were suggested to be a reaction to oxygen and sulfur radicals produced by the symbionts (Pruski and Dixon, 2007). Higher HSP70 levels were furthermore shown to concur with a higher stress tolerance in *Mytilus edulis* (Tedengren et al., 2000). While the exact function of USPs is still unknown, they have been observed to be expressed in *E. coli* under various stress conditions (Kvint et al., 2003). In metazoans, proteins with USP domains could have diverse functions in different taxa. USPs of *Hydra* might be involved in protection against microbial infection (Forêt et al., 2011). The *Riftia* USPs could therefore be involved in stress reactions, e.g. caused by symbiont digestion and subsequent host bacteriocyte death, and in interaction with the symbiont population.

Apoptosis-related proteins, specifically calpains and caspases, on the other hand, were less abundant in the trophosome and rather abundant in the plume: We observed about 4 to 7 times lower levels of calpains in the trophosome than in the plume and vestimentum, and caspases were hardly detected at all in the trophosome, but almost exclusively present in the plume. Apoptosis is probably of higher importance in the plume, as this organ is in direct contact with sulfide-rich water, whereas in the trophosome, another mechanism could be involved in cell death (Main Text). In sulfide-rich environments ROS are generated (Tapley et al., 1999), which – like sulfide – could cause cell damage, necessitating fast cell turnover. In the trophosome, on the other hand, cell turnover might involve a cathepsin-dependent mechanism (Main Text).

### 3) Release of organic compounds as a possible mode of nutrient transfer from symbiont to host

Besides symbiont digestion (Main Text), release of organic carbon by intact symbionts (“milking”) might also be a relevant mode of nutrient transfer from symbiont to host (Bright et al., 2000; Felbeck and Jarchow, 1998). Our calculation in host tissues and symbionts with Calis-p (Kleiner et al., 2018) showed $\delta^{13}C$ values of -6.7 to -8.3 in host tissues, whereas the symbionts where slightly lighter with a $\delta^{13}C$ of -9.8 in S-rich and -11.7 in S-depleted symbionts (Supp. Figure S2). $CO_2$ fixation via the Calvin cycle yields organic matter with lighter $\delta^{13}C$ values than organic carbon generated by the rTCA cycle (Hügler and Sievert, 2011; Pearson, 2010). The $\delta^{13}C$ values we calculated for *Riftia*, which correspond well to those reported in the literature (Fisher et al., 1990), might therefore indicate selective differential export or leakage of small organic compounds derived from the symbiont's rTCA cycle, rather than from the Calvin cycle. On the other hand, the accuracy of the SIF method is limited compared to the isotope ratio mass spectrometry (IRMS) gold standard (Kleiner et al., 2018). Additionally, we did not find a dedicated sugar transporter or organic acid exporter in the symbiont proteome and the dissimilar isotopic ratios might also be explained by less discriminating anaplerotic reactions in host tissues, e.g. catalyzed by phophoenolpyruvate carboxykinase (PEPCK), which we detected in all tissues.

Additional circumstantial evidence for milking in the *Riftia* symbiosis might be provided by our comparison of S-rich and S-depleted trophosomes, in which we observed quite dissimilar abundances of digestion enzymes. In S-depleted *Riftia* specimens, symbionts likely secrete less (or no) carbon compounds than in S-rich specimens, due to reduced $CO_2$ fixation. Therefore, the host probably needs to increase digestion pressure on the symbionts to cover its own basic nutritional demands. In S-rich specimens on the other hand, lower digestion rates might be required because the symbionts leak more organic material to the host. Moreover, it might be possible that the host preferentially digests symbionts with lower S content, to avoid high sulfur levels. A similar strategy of preferred digestion of S-depleted symbionts was suggested for the *Codakia orbicularis* symbiosis (Caro et al., 2007). Whether milking really is a relevant process in the *Riftia* symbiosis remains to be elucidated. Possibly, the *Riftia* host relies on a combination of symbiont digestion and milking, where the relative importance of both processes depends on the environmental conditions.

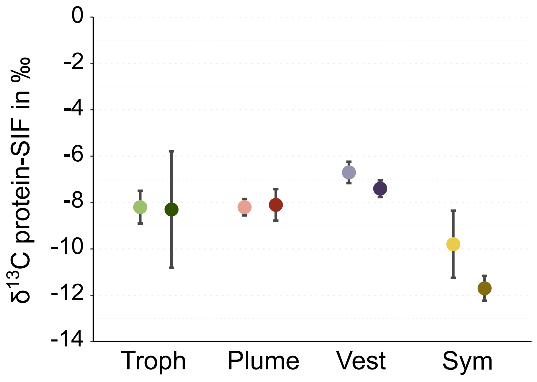


Supp. Figure S2: $\delta^{13}C$ values of *Riftia* host tissues and symbiont cells, calculated from the metaproteomics data using Calis-p (Kleiner et al., 2018). SIF: stable isotope fingerprinting. Light-colored symbols indicate S-rich tissues/symbionts, dark symbols indicate S-depleted tissues/symbionts. Error bars indicate standard error of the mean. Troph: trophosome, Vest: Vestimentum, Sym: enriched symbiont fraction.

### 4) Sulfur metabolism in host tissues

*Riftia* is well known to circumvent toxic effects of sulfide on its eukaryotic tissues by efficiently and reversibly binding sulfide to hemoglobins (Bailly and Vinogradov, 2005). In addition, the animal may be able to oxidize sulfide in its mitochondria, which would effectively also result in sulfide detoxification. We detected the mitochondrial host enzymes sulfide:quinone oxidoreductase, persulfide dioxygenase, thiosulfate sulfurtransferase and putative thiosulfate sulfurtransferases, which could serve to oxidize sulfide to the less toxic thiosulfate (Hildebrandt and Grieshaber, 2008). All these enzymes were low-abundant, however. In contrast to other animals in sulfidic habitats, for which mitochondrial sulfide oxidation is the only available mechanism to avoid sulfide toxicity (Grieshaber and Völkel, 1998; Powell and Somero, 1986), this strategy thus seems to be of minor importance in *Riftia*. For some thiotrophic symbionts, e.g. of *Bathymodiolus* mussels, thiosulfate appears to be the preferred sulfur form (Ponnudurai et al., 2017), and mitochondrial sulfide oxidation to thiosulfate thus likely also promotes symbiont energy generation. As *Riftia* symbionts prefer sulfide (even though they can probably also use thiosulfate; Markert et al., 2011; Robidart et al., 2011; Wilmot Jr. and Vetter, 1990), they would probably not benefit from mitochondrial sulfide oxidation by the host.

Sulfate, produced by the symbionts as end product of sulfide oxidation, could be excreted via the host's sodium-independent sulfate anion transporters detected in the trophosome (after being released from the symbionts with their own sulfate transporter). The host could even use this sulfate for its own biosynthetic processes via a bifunctional 3'-phosphoadenosine 5'-phosphosulfate synthase, which we detected in all vestimentum samples.

### 5) *Riftia* globins are even more diverse than previously thought

*Riftia's* extracellular hemoglobins have been thoroughly studied and are well known for their capacity to simultaneously bind $O_2$ and sulfide (Bailly and Vinogradov, 2005; Hourdez and Weber, 2005). In contrast, intracellular *Riftia* globins have received almost no attention thus far. Sanchez et al. (2007b) reported an intracellular *Riftia* globin, to which none of the globins detected in our study produces a close alignment. This indicates considerable diversity of this hitherto neglected protein group in *Riftia*. Intracellular globins have been described in few other annelid species; only three families are known to have circulating and non-circulating globins in addition to extracellular globins (reviewed in Bailly et al., 2007). *Riftia* might possess, in addition to its extracellular hemoglobins, also the other two globin types, which would make it the first described member of the *Siboglinidae* to have this trait: The related siboglinid worm *Lamellibrachia* has coelomocytes (van der Land and Nørrevang, 1977), indicating that these cells could also be present in *Riftia*. Coelomocytes could contain intracellular, circulating globins, while *Riftia's* intracellular neuroglobins might be non-circulating. Intra- and extracellular globins have also been described in the deep-sea alvinellid *Alvinella pompejana*, where they might buffer against variations in environmental $O_2$ concentration (Hourdez et al., 2000). In vertebrates, intracellular globins like cytoglobin and neuroglobin might be involved in intracellular $O_2$ storage and transport or may have a protective function against ROS or nitrous oxide. Cytoglobin might also have a role in collagen production (Hankeln et al., 2005). The role of *Riftia's* intracellular globins awaits further testing.

### 6) *Riftia* myohemerythrins as possible multi-purpose proteins

We detected myohemerythrins as additional possible $O_2$-binding molecules besides the well-described hemoglobins in *Riftia*. Hemerythrins are oxygen-binding molecules that can bind 15-28% more oxygen than simple hemes (Mangum, 1992). Myohemerythrin is hemerythrin localized in (annelid) muscle cells (reviewed in Vanin et al., 2006).

Our protein analysis showed a tissue-specific abundance pattern of the nine detected *Riftia* myohemerythrins. We therefore analyzed the sequences of these proteins to find indications for their respective functions. Additionally, we included hemerythrin sequences of annelids, a brachiopod and a member of the Bryozoa from the NCBI database (https://www.ncbi.nlm.nih.gov/) in our comparison (Supp. Figure S3, Supp. Table S11). Almost all of the conserved sites that exist in hemerythrins (as outlined by Stenkamp, 1994) are present in most *Riftia* myohemerythrins. One residue likely involved in $O_2$ binding has been described to either be a leucine or an isoleucine (Stenkamp, 1994). This residue corresponds to residue 76 (the positions refer to *Riftia*_myoHr6 in Supp. Figure S3), which is an isoleucine in most of the *Riftia* myohemerythrins, but a leucine in two sequences, both of which are significantly less abundant in the plume than in the trophosome samples. In the same sequences, residue 117 has been replaced by phenylalanine. This replacement has been noted before by Sanchez (2007) for one *Riftia* myohemerythrin sequence (ABW24415.1), which corresponds to Host_DN34237_c2_g1_i1::g.35857 analyzed in our study (Host_DN34237_c2_g1_i1::g.35857 is just 10 amino acids longer at the N terminus). Additionally, the six myohemerythrins with significantly higher abundance in the plume have several identical regions which differ from at least two of the other three myohemerythrins (Supp. Figure S3). While those are not assigned a specific function and we can therefore only speculate about their biological effect, these different variants nevertheless point to functional, tissue-specific diversification of the *Riftia* myohemerythrins. Possibly, the myohemerythrins in the trophosome are involved in interactions with the symbionts, whereas the plume-located myohemerythrins play a more general role in $O_2$ metabolism. Sanchez (2007) suggested that the trophosome-specific *Riftia* myohemerythrin they analyzed might be involved in protection against $O_2$ radicals, especially as potentially high radical production occurs in the trophosome. While reversible $O_2$ binding by an overexpressed *Riftia* myohemerythrin could not be shown by Sanchez (2007), this does not rule out *Riftia* myohemerythrin functioning as an $O_2$ carrier, as it is apparently not uncommon that purified hemerythrins do not bind $O_2$ (Costa-Paiva et al., 2017). Myohemerythrins have been noted to fulfill similar functions as myoglobin, which generate an $O_2$ gradient to the mitochondria (Coates and Decker, 2017). Heart and skeletal muscle cells as well as rhizobium nodules use myoglobin or leghemoglobin, respectively, to transport $O_2$ to the mitochondria/bacteroids. Leghemoglobin additionally likely protects the bacteroids against $O_2$ diffusion inside the cells, which contain the highly $O_2$-sensitive nitrogenase (reviewed in Wittenberg, 2007). *Riftia* myohemerythrin in the trophosome could fulfill a similar function in provisioning the symbionts with $O_2$ while protecting them from free $O_2$. This might also enable the symbionts to use the $O_2$-sensitive rTCA cycle while respiring $O_2$. Additionally, the *Riftia* myohemerythrin in the trophosome could be involved in regulating symbiont access to metals. For a hemerythrin-like protein in *Nereis diversicolor*, for example, it has been suggested that it regulates cadmium levels, as it binds cadmium in the gut (Demuynck et al., 2004).

*Riftia* myohemerythrins which are more abundant in the plume might be involved in maintaining an $O_2$ gradient across the plume and thereby facilitate oxygen diffusion into the animal. Additionally, they could store oxygen, as suggested for sipunculans during low tides (Mangum, 1992) and as proposed for the symbiotic oligochaete *O. algarvensis* (Wippler et al., 2016).

Moreover, *Riftia* myohemerythrins in general might serve further purposes as observed in other animals, e.g. iron storage, metal detoxification and functions in the immune system (Coates and Decker, 2017; Costa-Paiva et al., 2017; Sanchez, 2007; Seo et al., 2016). Potentially, they might even be involved in nitrate transport, as nitrate binding and concentrating by *Riftia* blood was observed (Hahlbeck et al., 2005). The exact function and (sub-) cellular localization of the different myohemerythrins deserves future studies, especially as an association with muscle cells in the non-muscular trophosome tissue seems unlikely. We rather speculate that myohemerythrin is contained in host bacteriocytes.

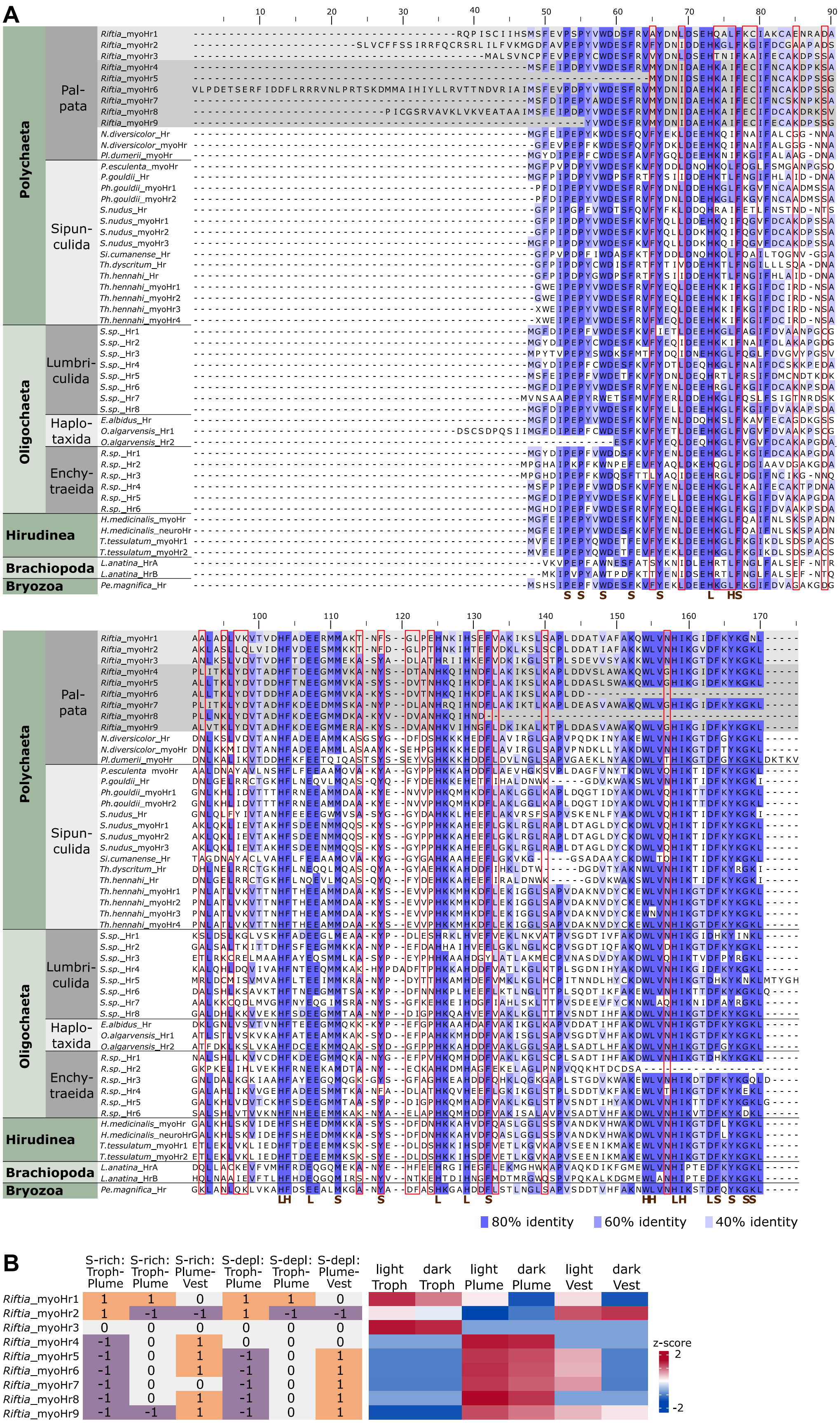
A
Polychaeta
Pal-pata
Sipun-culida
Oligochaeta
Lumbri-culida
Haplo-taxida
Enchy-traeida
Hirudinea
Brachiopoda
Bryozoa
80% identity
60% identity
40% identity
B
S-rich: Troph-Plume
S-rich: Troph-Plume
S-rich: Plume-Vest
S-depl: Troph-Plume
S-depl: Troph-Plume
S-depl: Plume-Vest
light Troph
dark Troph
light Plume
dark Plume
light Vest
dark Vest
Riftia_myoHr1 1 1 0 1 1 0
Riftia_myoHr2 1 -1 -1 1 -1 -1
Riftia_myoHr3 0 0 0 0 0 0
Riftia_myoHr4 -1 1 1 0 0 0
Riftia_myoHr5 -1 0 1 -1 0 1
Riftia_myoHr6 -1 0 1 -1 0 1
Riftia_myoHr7 -1 0 0 -1 0 1
Riftia_myoHr8 -1 0 1 -1 0 1
Riftia_myoHr9 -1 -1 1 -1 0 1
z-score
2
-2

221

Supp. Figure S3: A) Alignment of *Riftia* myohemerythrins with myohemerythrins of various other invertebrates (Supp. Table S11). Conserved residues according to Stenkamp (1994) are marked as follows: L: iron ligand binding, S: suggested to have a structural role, H: hydrophobic residues in $O_2$-binding pocket. Red boxes denote residues which are identical in one *Riftia* myohemerythrin subgroup (subgroups were separated by their distinct abundance patterns and are shown with different grey backgrounds) and different from at least two thirds of the sequences in the other subgroup. B) Left side, significance of abundance differences of detected myohemerythrins in *Riftia* tissues. 1: Significantly more abundant in the first tissue than in the second tissue, -1: Significantly less abundant in the first than second tissue, 0: no significant difference. Right side, heatmap of the z-scored %orgNSAF abundance of myohemerythrins in the different *Riftia* tissues. For accession numbers, see Supp. Table S11.

### 7) *Riftia* carbonic anhydrases are tissue-specific

*Riftia* expresses abundant carbonic anhydrases (CA) for interconversion of $CO_2$ and $HCO_3^-$ (Main text). Apparently, distinct isoforms dominate in different tissues (Main Text, Supp. Figure S4). Tissue-specific expression of *Riftia* CA has been noted before (De Cian et al., 2003b; Sanchez et al., 2007a). Three of the CAs detected in our study align closely to the CAs analyzed in these previous studies (Supp. Figure S4). Previous biochemical experiments hinted towards the existence of membrane-bound *Riftia* CA (De Cian et al., 2003a, 2003b). Indeed, predicted transmembrane helices and/or signal peptides in three of the CA sequences in our study indicate that these could be membrane-bound (or even secreted) rather than cytoplasmic. These three CAs also align more closely to each other than to the other CA sequences (Supp. Figure S4). Interestingly, putative membrane-bound CA isoforms seem not to be restricted to one tissue, but might increase $CO_2$ transport efficiency across different tissues. The residues involved in zinc binding and several of the residues forming the hydrogen bond of the active site according to De Cian et al. (2003b) seem not to be conserved across all *Riftia* CA isoforms, indicating possible functional diversification.

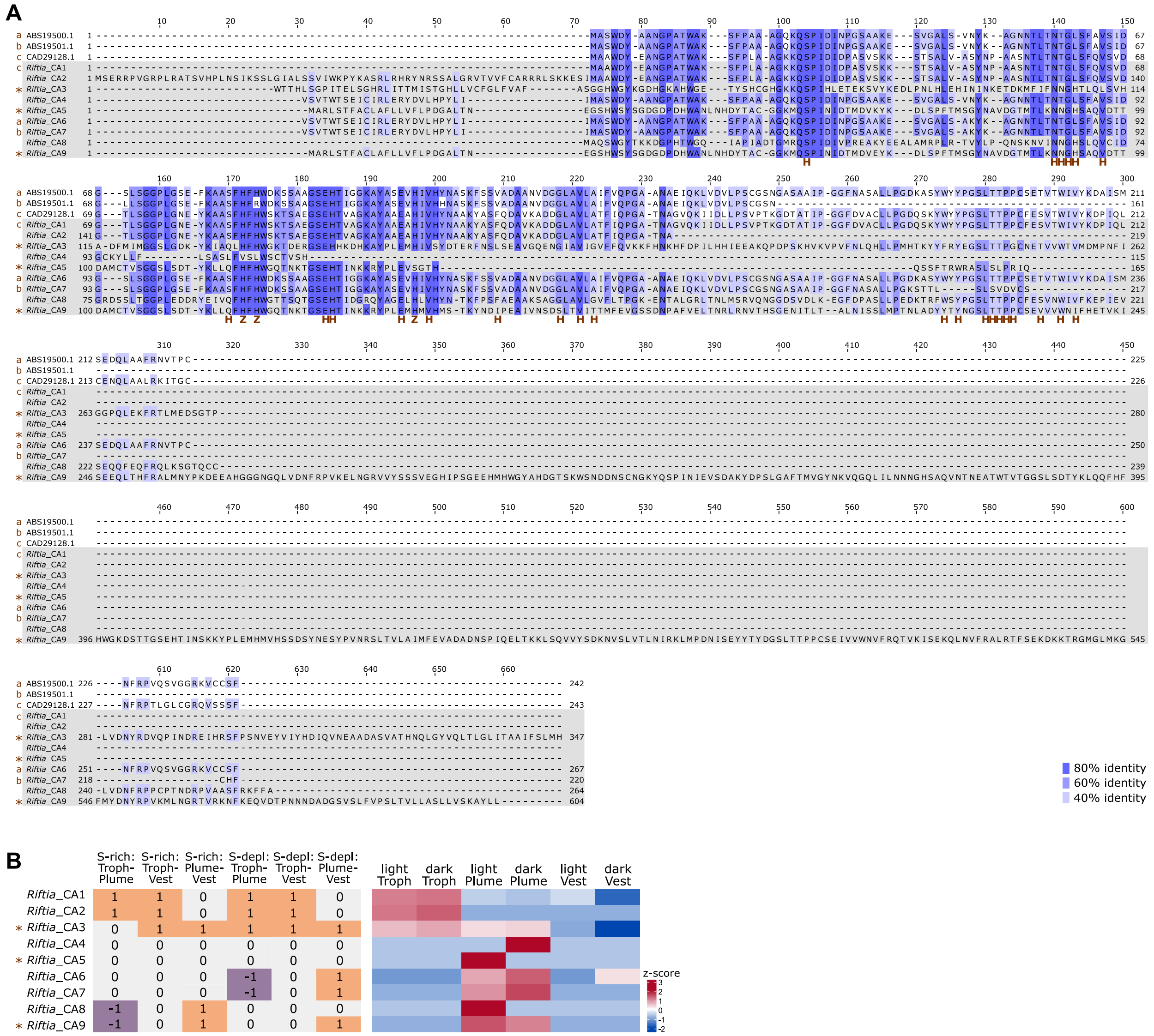


Supp. Figure S4: A) Alignment of carbonic anhydrases (CAs) detected in this study (Riftia_CA1 – Riftia_CA9) and CAs detected in previous studies (De Cian et al., 2003b; Sanchez et al., 2007a), for accession numbers see Supp. Table S12). Lower case letters indicate which CA sequences identified in our study align most closely to the sequences identified in previous studies. Asterisks mark potentially membrane-bound/secreted CA isoforms. Capital letters mark residues involved in functions according to De Cian et al. (2003b). H: residues forming the hydrogen bond network of the active site, Z: histidine residues involved in zinc binding. B) Left side, significance of abundance differences of detected CAs in the *Riftia* tissue comparisons. 1: Significantly more abundant in the first tissue than in the second tissue, -1: Significantly less abundant in the first than second tissue, 0: no significant difference. Right side, heatmap of the z-scored %orgNSAF abundance of CA in the different *Riftia* tissues. For accession numbers, see Supp. Table S12.

### 8) $C_4$ metabolism in the *Riftia* host?

We detected proteins involved in transport of inorganic carbon by the *Riftia* host to its symbiont, i.e., V-type ATPase subunits, carbonic anhydrase and bicarbonate exchangers (Main Text Figure 2). In addition, it has previously been suggested that the host also provides the symbionts with pre-fixed $CO_2$, i.e. with small organic compounds (Felbeck, 1985). Phosphoenolpyruvate carboxykinase, which we detected in all host tissues, and pyruvate carboxylase, which was only lacking in S-rich plumes, catalyze incorporation of $CO_2$ into oxaloacetate, which would then be available for host and symbiont metabolism. Like for the *Riftia* symbiosis, host pre-concentration of $CO_2$ via phosphoenolpyruvate carboxykinase has been postulated for the *B. azoricus* symbiosis (Ponnudurai et al., 2017). On the other hand, pre-fixation of $CO_2$ in the *Riftia* symbiosis has been doubted by Childress et al. (1984), who observed high levels of total $CO_2$ and $CO_2$ partial pressure in the blood of the tubeworm. Following this, $^{14}C$-labelled malate and succinate formation in the plume observed by Felbeck (1985) could also be due to anaplerotic host metabolism, rather than presenting a means to sustain the symbionts. If host prefixation of $CO_2$ would occur, the symbiont would either need transporters for uptake of the organic compounds, or the organic compounds would needed to be decarboxylated inside the host cells, so that the symbiont could take up $CO_2$. We found five symbiont TRAP transporter components in the trophosome samples, two of which were predicted to transport C4 dicarboxylates, which could take up small organic $CO_2$ carriers. At the same time, import of these organic $CO_2$ transport molecules into the bacteriocyte (and maybe the symbiont cell) would probably rule out simultaneous leakage or export of small organic molecules by the symbiont to sustain the host (milking, see above), as uptake and concomitant release of small organic molecules by the bacteriocyte seems unlikely. Therefore, it remains speculative whether the *Riftia* host pre-fixes $CO_2$, also in light of our metaproteomics data.

### 9) Nitrogen metabolism in the *Riftia* symbiosis

Our reconstruction of nitrogen metabolic pathways in the *Riftia* host and symbiont revealed that *Riftia* depends on ammonium supplied by the symbiont, but can synthesize pyrimidines and polyamines independently (main text, detailed below), while the host might degrade nitrogen-containing symbiont waste products.

We found two possible $NO_3^-$ transporters (probable peptide/nitrate transporter At3g43790: Host_DN35983_c1_g1_i7::g.173229, Host_DN35983_c1_g1_i4::g.173222) in the *Riftia* transcriptomes of all tissues, although not in the metaproteome. The respective proteins could be involved in $NO_3^-$ uptake from the environment and $NO_3^-$ transport inside the animal. Hypothetically, *Riftia* myohemerythrin, which was abundantly detected in the host metaproteome, might serve as a nitrate-binding component in the blood, in addition to hemoglobin, which has previously been suggested to bind and concentrate $NO_3^-$ (Hahlbeck et al., 2005). After transport to the symbionts, nitrate could be ammonified via the symbionts' membrane-bound nitrate reductase (NarGHIJ) and nitrite reductase (NirBD) (Main Text Figure 4). Although NarGHIJ is mainly known to be involved in dissimilatory nitrate reduction, i.e., respiration, assimilatory nitrate reduction via membrane-bound NarGHJI and NirBD was demonstrated in *Mycobacterium tuberculosis* (Malm et al., 2009) and suggested for the symbiont of the tubeworm *Ridgeia piscesae* (Liao et al., 2014). Symbiont periplasmic nitrate reductase NapAB, of which we detected NapB in the metaproteome, could theoretically also be involved in assimilatory nitrate reduction (Liao et al., 2014, reviewed in Sparacino-Watkins et al., 2014). Both partners of the *Riftia* symbiosis could then incorporate ammonium into their organic matter via glutamine synthetase/glutamate synthase (GS-GOGAT cycle) as well as via glutamate dehydrogenase, all of which we detected on the protein level in host and symbiont. Host glutamate dehydrogenase was overall most abundant in the trophosome, compared to other tissues, possibly because the host metabolizes ammonia generated by the symbionts. Eukaryotic ammonium assimilation via the GS-GOGAT cycle and via glutamate dehydrogenase has previously been suggested in silkworms (Hirayama et al., 1997, 1998), mosquitoes (Scaraffia et al., 2005) and *R. piscesae* tubeworms (Liao et al., 2014).

Both symbiotic partners appear to be capable of producing the polyamine putrescine (Main Text Figure 4). This contradicts a previous, enzyme activity-based study, in which arginine decarboxylase and ornithine decarboxylase activity were presumed to be exclusively due to symbiont enzymes (Minic and Hervé, 2003). Given the ubiquitous importance of polyamines (see below), it is probably advantageous for the host to be less dependent on the symbiont for polyamine synthesis, even more so during times of reduced symbiont digestion (i.e. in S-rich specimens, Main Text).

Our results suggest that *Riftia* degrades nitrogenous compounds via the urea cycle. We detected all urea cycle enzymes (except for nitroxide synthase and arginine deaminase, which are not required for the full cycle) in the host's proteome, although not all proteins were detected in all tissues. We also identified a urea-proton symporter, probably involved in urea excretion, in all plume samples and one vestimentum sample. Interestingly, the symbiont, unlike the host, does not seem to possess a complete urea cycle (Main Text Figure 4). As host arginase was significantly more abundant in the trophosome than in the other tissues, it probably serves to break down nitrogen-containing metabolites of host and symbiont. Taken together, our results point to a complete urea cycle in the *Riftia* trophosome, as previously indicated by biochemical analyses (De Cian et al., 2000).

### 10) Host polyamines may be involved in host-symbiont interactions

We detected host spermine synthase and spermidine synthase, which synthesize the polyamines spermine and spermidine, only in the trophosome metaproteomes, but not in other tissues. Therefore, *Riftia* polyamines could have a role in symbiont-host interactions. Such a role for a polyamine has been suggested for *Shigella flexneri*, where cadaverine is likely involved in restricting the pathogen to the host phagocytic vacuole (Fernandez et al., 2001). Similarly, *Riftia* host polyamines could be involved in keeping the symbiont from spreading in the host tissue. Additionally, spermine could play a role in immune system suppression in the trophosome, as it has been shown to suppress an innate inflammation reaction in human cell cultures after lipopolysaccharide stimulation (Zhang et al., 1999). In addition to these putative interaction-related functions, the polyamines of both *Riftia* host and symbiont probably also have more general roles: Polyamines are known to interact with DNA, RNA and proteins, are involved in reaction to oxidative and acid stress and apoptosis, and play a role in bacterial outer membrane biogenesis and functions. Polyamine levels are higher in actively proliferating cells (reviewed in Igarashi and Kashiwagi, 2010; Shah and Swiatlo, 2008; Wallace et al., 2003). This may explain the high abundance of polyamine-generating enzymes in *Riftia* trophosome, where host and symbiont cells proliferate at very high rates in the central parts of the trophosome lobules (Pflugfelder et al., 2009).

### 11) High FIH abundance points to hypoxic conditions in the *Riftia* trophosome

Our results indicate that *Riftia* sustains hypoxic conditions in the trophosome, which would benefit the microaerophilic symbionts. The hypoxia-inducible factor 1-alpha inhibitors (factor inhibiting HIF1a; FIH), which we detected almost exclusively in trophosome samples, are possibly involved in metabolic regulation under these conditions. FIH inhibits HIF1a, which is a main regulator of hypoxia-dependent transcription regulation, during normoxic conditions (Ferguson III et al., 2007). While we did not detect HIF1a on the protein level, the presence of FIH very likely indicates that HIF1a is also expressed. A low (but detectable) abundance of FIH might indicate that HIF1a is not inhibited and the trophosome is hypoxic. High FIH abundance, on the other hand, would indicate HIF1a inhibition and thus more normoxic conditions, e.g. during temporary phases of higher $O_2$ concentration. The hypothesis of a hypoxic trophosome is in line with the observation that *Caenorhabditis elegans* HIF1a is induced by hypoxia and rapidly degraded under normoxic conditions

(Shen and Powell-Coffman, 2003). Further studies are needed to verify the presence of HIF1a and to elucidate the regulation of FIH itself under different $O_2$ regimes in *Riftia* trophosomes.

### 12) The *Riftia* immune system might protect the symbiosis against phages

The symbionts appear not to elicit a general host immune response, as a number of host immune proteins such as lysozyme, tyrosine-protein kinases and peptidoglycan recognition proteins were detected, but overall, their abundance in the trophosome was not higher than in other tissues (Main Text Figure 2). Galaxins, proteins associated with coral exoskeletons (Watanabe et al., 2003), have been implied in immune system and symbiont interaction functions in squids (Heath-Heckman et al., 2014). While one galaxin transcript was detected before in the *Riftia* body wall (Sanchez et al., 2007b), we only detected galaxins in the firm vestimentum tissue, implying that they could have a structural rather than an immune system function in *Riftia*. Most likely, the host vesicles in which the symbionts are contained shields the bacteria against recognition and attack by the *Riftia* immune system. Similarly, in chronic infections of insect hosts with endosymbionts, the host's immune system does not eliminate the symbionts (Ratzka et al., 2012). Rather than acting against the symbionts, the *Riftia* immune system probably protects the host (and thereby the symbiosis as a whole) against infections, and might even be involved in protecting the symbionts against phage infections. The symbiont population is basically a bacterial monoculture inside *Riftia*, for which phage infections could easily be fatal. The detection of CRISPR-Cas-associated proteins in the *Riftia* symbiont metaproteome might indicate that phage infections play a role during symbiosis. Symbionts could be prone to infection also inside the trophosome, as phages can rapidly penetrate tissues of higher organisms (Dabrowska et al., 2005). Therefore, host proteins likely involved in interaction with viruses, e.g. interferon-induced GTP-binding proteins, glycylpeptide N-tetradecanoyltransferase 2 and deoxynucleoside triphosphate triphosphohydrolase SAMHD1, could not only protect *Riftia* itself against a viral infection, but could putatively also be involved in shielding the symbionts against phages. The *Riftia* immune system could also be involved in initial identification of the symbiont during host colonization (e.g. via lectins which are present in the host's proteome). In the symbiotic stage, however, few and specific functions of the immune system seem to be harnessed for symbiotic interactions.

### 13) Roles of eukaryote-like proteins in *Ca.* E. persephone and other host-associated and free-living organisms

We compared the proportion of proteins with putative roles in host-symbiont interactions, especially eukaryote-like proteins (ELPs), which are encoded in the metagenomes of several (endo)symbionts, pathogens, free-living bacteria and environmental samples (Main Text Figure 5, Supp Figure S5, Supp. Table S4, Supp. Table S5). The three *Riftia*/*Tevnia* symbiont metagenomes that were included in our protein database (see methods) showed, as expected, almost identical distribution patterns of the respective protein groups. As *Riftia* and *Tevnia jerichonana* are colonized by the same symbiont species (Gardebrecht et al., 2012; Perez and Juniper, 2016), differences are probably due to unequal sequencing coverage. The pattern observed in the *Ridgeia* symbiont metagenomes was very similar to that in the *Riftia/Tevnia* symbiont, which is in line with the fact that the *Ridgeia* symbiont likely belongs to the same species as the *Riftia* and *Tevnia* symbiont (Perez and Juniper, 2016). The similarity of ELP patterns between the tubeworm symbionts and symbionts of shallow-water bivalves (Supp. Figure S5) suggests parallels in interaction strategies with their respective hosts.

We also observed differences in ELP distribution patterns between distinct organism groups (Supp. Figure S5), which points to specific environmental adaptations. Armadillo repeats, FG-GAP, GIDE, Hyr, Integrins, and TSP domains were almost exclusively found in the gammaproteobacterial sulfur-oxidizing endosymbionts, suggesting specialized roles in these associations. The low number of the respective genes in the *B. azoricus* sulfur-oxidizing endosymbiont metagenomes could be an artifact caused by the small and presumably rather incomplete metagenome databases available. On the other hand, whereas *Riftia* endosymbionts are probably actively digested by the host – and even more so in times of starvation – in *B. azoricus*, autolysis of senescent bacteria seems to be part of the normal cell cycle (Kádár et al., 2008). This could explain a higher abundance of putative host-interaction-related and presumably phagocytosis-inhibiting proteins in the *Riftia* symbiont, and in the symbionts of the shallow-water bivalves, which are probably also actively digested by their hosts (Caro et al., 2009; Johnson and Fernandez, 2001; Stewart and Cavanaugh, 2006). The *B. thermophilus* endosymbiont ELP distribution pattern was rather distinct from that of all other gammaproteobacterial SOX endosymbionts. The reasons for these differences, including especially the high abundance of attachment-related proteins in the *B. thermophilus* endosymbiont, remain open for debate (Ponnudurai et al., in rev.).

Some domains, especially ankyrins, PD40 and WD40, Sel1, SNARE, TPR and coiled-coil domains are rather widespread and are also found in non-host-associated bacteria. This points to more general functions, which might have been harnessed in the course of symbiotic adaptations. Some of the analyzed host-associated bacteria, including the *Riftia* symbiont, are horizontally transmitted, i.e. taken up from the environment (Bright and Bulgheresi, 2010; Nussbaumer et al., 2006). This could explain the relatively high similarity of these bacteria to the protein group pattern of free-living organisms and the diversity of protein groups. In comparison, obligate endosymbionts (including *Calyptogena* endosymbionts, Kuwahara et al., 2007), which undergo genome reduction and genetic drift, showed a reduced diversity of the analyzed protein groups. Nipsnap domains are mainly found in rhizobia and could thus be involved more specifically in rhizobia-plant interactions. As expected, the vent- and chimney metagenomes, which encompass a variety of prokaryotic and eukaryotic microorganisms, had the highest total number of eukaryote-like domain-containing proteins. Nevertheless, relative to the total number of proteins in the respective (meta)genomes, some of these protein groups are under-represented in comparison to the single bacteria analyzed, which indicates that ELP patterns may vary substantially between microbes and appear to have specific functions in individual bacterial groups.

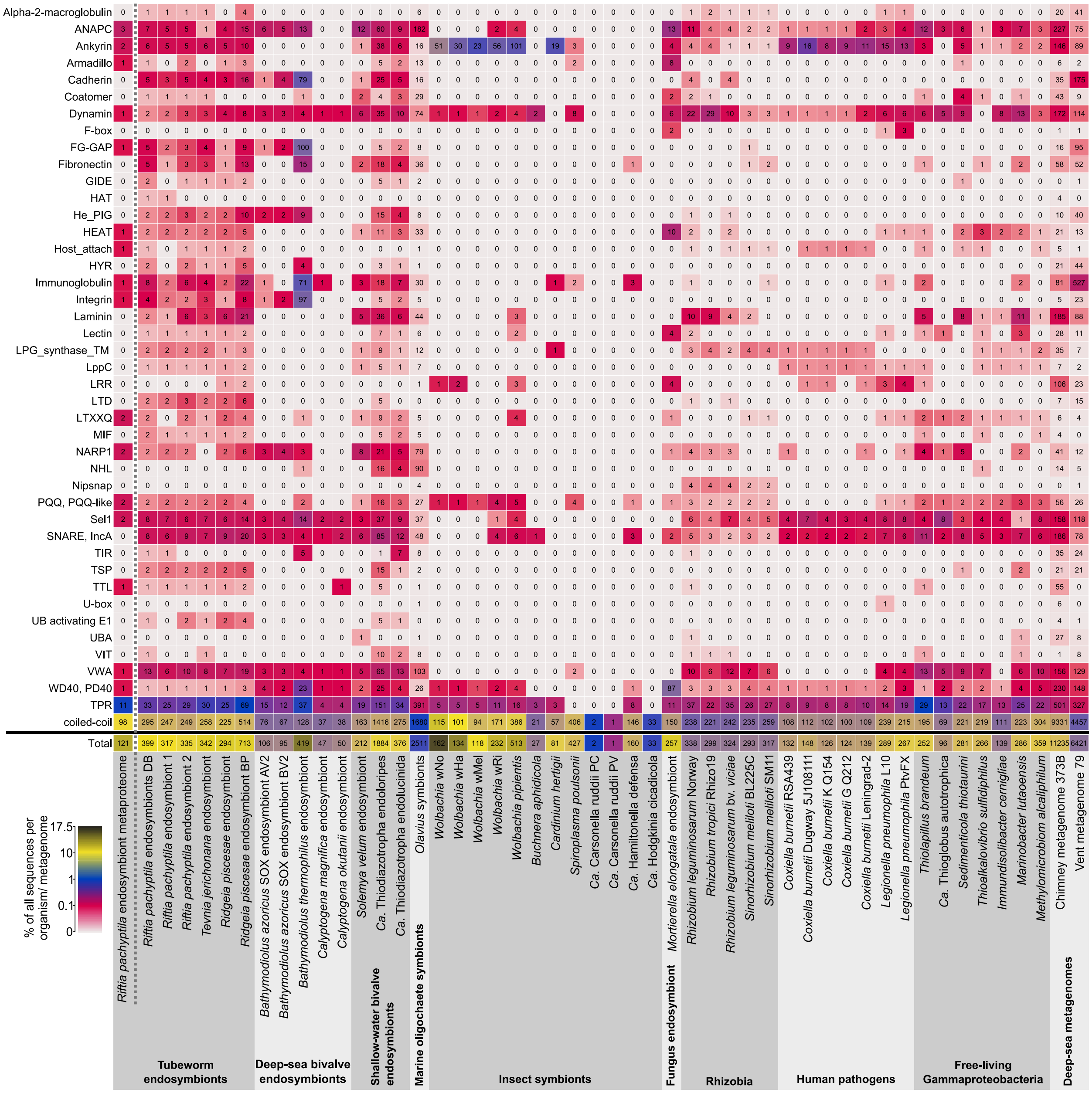


Supp. Figure S5: Selected domains with eukaryote-like structures and with putative functions in symbiont-host interactions in the *Riftia* symbiont and in selected other organism groups and environmental metagenomes. Color scale: percentage of genes/proteins with the respective domain relative to all genes/proteins, numbers: total number of genes/proteins with the respective domain. For details on the organisms, see Supp. Table S5. For further information about the selected protein groups, see Supp. Table S4. '*Riftia pachyptila* endosymbiont metaproteome' refers to *Riftia* symbiont proteins detected in this study.

### 14) Symbiosis-relevant proteins of *Ca.* E. persephone may facilitate host interactions

**Symbiont proteins might modulate host gene expression.** Our metaproteomic analyses suggest that host histones could interfere with the symbiont, because they can be precursors for antimicrobial peptides (AMPs, Main Text). The symbiont, on the other hand, could itself manipulate the hosts' histones, and thereby the host transcription regulation: We found the SET-domain containing protein Sym_EGV51182.1 in the *Riftia* symbionts' metagenome (albeit not on the protein level). Sym_EGV51182.1 had highest sequence similarity to several eukaryotic histone-lysine N-methyltransferases with 30-32% identity (blastp against Uniprot/Swissprot, https://www.expasy.org/). In *L. pneumophila*, a lowly expressed SET domain-containing methyltransferase trimethylates host histone H3 and thereby probably represses the expression of host genes, including genes of the innate immune system (Rolando et al., 2013). The *Riftia* symbiont SET domain-containing protein might have a similar function and modulate gene expression of the host, possibly enabling symbiont persistence by repressing the host's immune response.

***Ca.* E. persephone dodecin and porin may facilitate symbiont survival.** We detected a putative symbiont dodecin (Sym_2601633487) in about 30 times higher abundance in symbionts from S-depleted trophosomes (0.886%orgNSAF, rank 16) than in symbionts from S-rich trophosomes (0.028%orgNSAF, rank 409). This protein could be involved in adaptation to conditions under S depletion, e.g. to a putatively increased digestion pressure. A blast search of the *Ca.* E. persephone putative dodecin against the Uniprot/Swissprot database (https://web.expasy.org/blast/) showed highest similarity to a *Halorhodospira halophila* dodecin (54% identity), followed by a *Thermus thermophilus* dodecin (45% identity). *H. halophila* dodecin was shown to bind the flavin FMN (Grininger et al., 2009). Likewise, the dodecin of *T. thermophilus* binds FMN and coenzyme A. Dodecin could therefore work as a flavin trap, for example when flavins are released from denatured flavoproteins (Meissner et al., 2007). Reduced flavins and flavoproteins generate ROS (Massey,

1994). The *Ca.* E. persephone dodecin could therefore counteract the detrimental effects of putatively higher oxygen levels in S-depleted trophosomes by sequestering flavins, especially as probably less flavins are needed for redox reactions and electron transfers and additionally flavoproteins might be degraded due to the energy-limited conditions. Higher stress levels due to digestion could also facilitate flavin release from symbiont proteins. Sequestering these flavins by dodecin would decrease the potential for ROS formation. Specific involvement of dodecin in intracellular survival has been speculated for *M. tuberculosis*. This protein could be involved in survival of the pathogen under the acidic conditions in lysosomes, as the FMN:dodecin complex is more stable at a lower pH (Bourdeaux et al., 2018). Likewise, the *Riftia* dodecin could exert a protective influence during increased symbiont digestion in S-depleted trophosomes.

Additionally, symbiont porins were the most abundant proteins (9.8%orgNSAF) in S-depleted symbionts and about four times more abundant than in S-rich symbionts. Porins could be involved in interactions with the host by exporting symbiont effectors, or by im- or exporting host effectors, e.g. to modulate digestion (see also Main Text). Their higher abundance in S-depleted trophosomes could therefore be a reaction to the postulated higher digestion pressure. In the genetic neighborhood of the *Riftia* symbiont porin (Riftia1 genome) are several genes with predicted functions in lipopolysaccharide biosynthesis, and sugar metabolism and RNA metabolism. The exact function of the *Ca.* E. persephone porin remains to be elucidated, however.

***Ca*****. E. persephone might be naturally competent during symbiosis.** Several of the symbiont proteins detected in this study indicate that the *Riftia* symbiont population is genetically competent during symbiosis and might exchange genetic material. The type 4 pilus (T4P) system might be involved in natural transformation (Main Text). Besides the T4P system, we identified proteins belonging to toxin-antitoxin-systems (putative zeta toxins Sym_EGV52816.1, Sym_EGW56021.1).

These can be encoded on conjugative elements, so that bacteria which do not have these conjugative elements are killed (Wozniak and Waldor, 2009). Additionally, proteins of the Tol-Pal-system were present in the *Ca*. E. persephone metaproteome. This system not only confers outer membrane stability, but is also involved in uptake of the mostly plasmid-encoded bacteriocin colicin A in *E. coli* (Lazzaroni et al., 2002; Yang et al., 2014). Moreover, we detected two plasmid-related proteins (Sym_EGV50018.1 – plasmid maintenance system killer, Sym_EGV50285.1 - TraB family pAD1 protein). Previous genome analyses showed that the *Riftia* symbiont has an integrated partial F plasmid (Riftia1 genome; Gardebrecht et al., 2012). Conjugation could allow the symbionts to exchange genetic material, potentially with a role in symbiosis. These results indicate that strain adaptation and diversification of the *Riftia* symbiont might take place under symbiotic conditions.

### 15) Metabolite exchange between the symbiosis and the environment depends on S availability

Lower S availability seems to not only impact the *Riftia* host's nutrition, but also its exchange of compounds with the environment and thereby its capacity to support the symbiont: We detected carbonic anhydrases (CAs) and FIH in higher total abundance in S-depleted specimens (summed up %orgNSAF), whereas V-type ATPase subunits, myohemerythrin and cytoplasmic malate dehydrogenase were overall less abundant in S-depleted specimens. These differences are probably due to changes in the habitat chemistry: Sites with low sulfide concentrations (leading to low S availability for the symbiosis) are alkaline, whereas those with higher sulfide levels have an acidic to neutral pH (Scott et al., 2012). At the same time, higher sulfide concentrations implicate higher $CO_2$ levels, but lower $O_2$ concentrations. In plumes of *Riftia* specimens from habitats with low sulfide-levels (and thus also low $CO_2$ levels), higher activities of carbonic anhydrase where measured (Scott et al., 2012). The higher abundances of carbonic anhydrases in trophosomes and plumes of S-depleted specimens thus probably serve to enable $CO_2$ uptake against a shallower concentration gradient (if $CO_2$ is taken up at all under these conditions). Moreover, as autotrophic metabolism of the symbionts is probably inhibited during sulfide depletion (Childress et al., 1991), $CO_2$ generated by host and symbiont respiration would need to be excreted in order to prevent metabolic acidosis. CA could then function to catalyze the conversion of bicarbonate to $CO_2$ to facilitate $CO_2$ diffusion into the environment. At the same time, the need for pH regulation seems to be decreased in S-depleted *Riftia*, as we detected a lower sum abundance of V-type ATPase subunits in plumes of these specimens. In S-depleted worms, less protons are generated during symbiont sulfide oxidation, which diminishes the need for ATP-consuming proton excretion.

Whereas reduced sulfur compounds, and potentially $CO_2$, are probably limiting under S-depleted conditions, $O_2$ could be limiting in S-rich specimens: Myohemerythrin, which is potentially involved in $O_2$ storage and transport, had an about 2.9 times higher overall abundance in S-rich plumes and trophosomes than in S-depleted plumes and trophosomes. It might facilitate $O_2$ uptake and $O_2$ provision of the symbionts, which are relatively more abundant, and thus in total likely need more $O_2$, in S-rich trophosomes. At the same time, trophosomes of S-rich specimens apparently still have lower free $O_2$ levels, as indicated by about 10 times lower levels of FIH in S-rich trophosomes and higher malate dehydrogenase abundance (see above and Main Text). Although on the one hand, hypoxic conditions would be beneficial for the microaerophilic symbionts, on the other hand, too low $O_2$ concentrations could limit sulfur oxidation and add to elemental sulfur accumulation in symbionts of S-rich *Riftia* specimens. While the symbionts are possibly capable of $NO_3^-$ respiration, this pathway probably plays only a minor role compared to $O_2$ respiration (Girguis et al., 2000). Additionally, the energy gained from $NO_3^-$ respiration would be lower than that from aerobic respiration, due to the lower redox potential of $NO_3^-$. The higher myohemerythrin abundance thus likely points to an increased $O_2$ demand in S-rich *Riftia* specimens. Similarly, myoglobin concentrations in vertebrate muscles increase with oxygen demand, and myoglobin expression depends at least partially on $O_2$ availability (reviewed in Wittenberg, 2007).

The symbiont metaproteome profile also hints at more oxic conditions in S-depleted than S-rich trophosome: Two of the symbiont antioxidant proteins (Sym_EGV49864.1 – rubrerythrin, Sym_EGV52457.1 – putative hemerythrin) were significantly more abundant in the S-depleted symbionts. Moreover, it could be speculated that the symbionts are killed before digestion via an oxidative burst, as is the case for intracellular pathogenic bacteria (Schramm et al., 2014). The higher antioxidant levels in S-depleted symbionts would then be a means to counteract the host-produced ROS. A role of hemerythrin for survival inside host macrophages via ROS detoxification has been suggested for *Aeromonas hydrophila*, a gram-negative gammaproteobacterium (Zeng et al., 2016).
