## Supplementary Table S4 for "Host-microbe interactions in the chemosynthetic *Riftia pachyptila* symbiosis"

Supp. Table S4: Domains and protein families with a putative role in host-symbiont interactions. The domains and protein families listed here were included in the comparisons in Figure 5 and Supp. Figure S5, which show the percentage of the respective protein groups in the *Riftia* symbiont metagenome and in metagenomes of other symbiotic and free-living organisms. % bacterial, total number bacterial: Percentage and total number of bacterial species in which this domain is found in the SMART database (January 2019).

| **Domain name** | **Pfam/SMART annotation** | **% bacterial (total number bacterial)** | **Literature/comment** |
| --- | --- | --- | --- |
| Alpha-2-macroglobulin | alpha-2-macroglobulin family (A2M), including N-terminal MG1 domain | A2M: 42.05% (2057) | A2Ms: protease inhibitors which are important for eukaryotic innate immunity, if present in prokaryotes apparently fulfill a similar role, e.g. protection against host proteases (Wong and Dessen, 2014) |
| ANAPC | Anaphase-promoting complex subunits | APC2: 0 | Ubiquitin ligase, important for cell cycle control in eukaryotes (Peters, 2006)  Bacterial proteins might interact with ubiquitination pathways in the host (Rytkönen and Holden, 2007) |
| Ankyrin | Ankyrin repeats | 10.88% (8348) | Mediate protein-protein interactions without sequence specificity (Li et al., 2006)  Sponge symbiont ankyrin-repeat proteins inhibit amoebal phagocytosis (Nguyen et al., 2014)  Present in sponge microbiome metatranscriptomes, putative role in symbiont-host interactions (Díez-Vives et al., 2017)  Present in obligate intracellular amoeba symbiont *Candidatus* Amoebophilus asiaticus genome, probable function in interactions with the host (Schmitz-Esser et al., 2010) |
| Armadillo | Armadillo repeats | 0.83 % (67) | Eukaryotic armadillo repeats are involved in protein-protein interactions, e.g. in intracellular signaling and cytoskeletal organization (Coates, 2003) |
| Cadherin | Cadherin domains, Cadherin-homologous | CA: 6.4% (956)  CADG: 47.77% (739) | Calcium-dependent cell-cell adhesion, tissue morphogenesis in animals (Koch et al., 2004)  Present in sponge microbiome metatranscriptomes, putative role in symbiont-host interactions (Díez-Vives et al., 2017)  Putative mediation of protein interactions and cell-cell adhesion in the marine bacterium *Saccharophagus degradans*; most cadherin-containing prokaryotes are aquatic (Fraiberg et al., 2010) |
| Coatomer | coatomer complex units | NA | Formation of vesicles for intracellular transport (Wang et al., 2016) |
| coiled-coil | coiled-coil domains | NA | Structural domain in all three domains of life, in proteins of diverse functions (Truebestein and Leonard, 2016)  Virulence effectors secreted by type III secretion systems of pathogenic bacteria often contain coiled-coil domains, which e.g. could interact with host signaling pathways (Delahay and Frankel, 2002) |
| Dynamin | Dynamin family | DYNc: 0 | Large GTPases which in eukaryotes are important for vesicle scission and lipid tabulation and fission (Praefcke and McMahon, 2004)  Conserved in many bacterial genomes, cellular role of bacterial dynamins not clear, could be involved in cytokinesis under osmotic stress (Bramkamp, 2012) |
| F-box | ubiquitin interacting | 0.42% (92) | Present in obligate intracellular amoeba symbiont *Candidatus* Amoebophilus asiaticus genome, could interact with host ubiquitination pathways (Schmitz-Esser et al., 2010)  Bacterial proteins might interact with ubiquitination pathways in the host (Rytkönen and Holden, 2007) |
| FG-GAP | Extracellular repeat in alpha integrins | NA | *Leptospira* FG-GAP proteins could be involved in interaction with host tissues (Chou et al., 2012)  Also see Integrin |
| Fibronectin | Fibronectin (FN) and FN-like | FN1: 0  FN2: 0  FN3: 24.19% (11706) | Present in sponge microbiome metatranscriptomes, putative role in symbiont-host interactions (Díez-Vives et al., 2017) |
| GIDE | E3 ubiquitin ligase | NA | Bacterial proteins might interact with ubiquitin pathways in the host (Rytkönen and Holden, 2007) |
| HAT | Half A TPR repeat | 2.00% (86) | RNA and peptide binding motif, involved in RNA metabolism, related to TPR repeat, highly conserved in eukaryotes, almost all HAT proteins are found in nucleus (Hammani et al., 2012) |
| He_PIG | putative immunoglobulin | NA | See Immunoglobulin |
| HEAT | HEAT and HEAT-like repeats | NA | Found in eukaryotic proteins with different functions, involved in protein-protein interactions, related to armadillo and ankyrin repeats (Yoshimura and Hirano, 2016) |
| Host_attach | bacterial attachment to host cells | NA | Required for attachment to host cells (Matthysse et al., 2000) |
| HYR | hyalin repeat domain | NA | Belongs to immunoglobulin-like fold, probably involved in cell adhesion in eukaryotes (Callebaut et al., 2000) |
| Immuno-globulin | Immunoglobulin, immunoglobulin-like | 1.24% (624) | Present in functionally diverse proteins, involved in molecular recognition, in pathogenic *E. coli* strains important for host cell infection, components of adhesins and other proteins (Bodelón et al., 2013)  Bacterial surface proteins could be involved in recognition between symbionts and host |
| Integrin | Integrin alpha and beta subunits | Int_alpha: 26.37% (1071)  INB: 1.2% (16) | Integrins are important for cell adhesion and signaling in metazoans, also present in apusozoans; uncommon in prokaryotes (Sebé-Pedrós et al., 2010)  Bacterial structures not found to be full-length integrins, but full-length beta-propeller domains are present, function unknown (Chouhan et al., 2011)  Could be involved in interaction with host tissues in *Leptospira* (Chou et al., 2012) |
| Laminin | Laminin domains | LamNT: 0.84% (22)  LamB: 0  LamG: 6.49% (544)  EGF_Lam: 0 | Important constituent of basement membranes in animals (Sasaki et al., 2004) |
| Lectin | lectins | Jacalin: 7.49% (96)  Gal-bind_lectin: 0.08% (2)  B_lectin: 5.58% (219)  CLECT: 1.54% (263)  GLECT: 0.13% (3) | Carbohydrate-binding proteins, involved in immune system reaction by pathogen recognition and aggregation in vertebrates and invertebrates (Fujita, 2002)  Virulence factors in pathogens (Imberty et al., 2004) |
| LPG_synthase_TM | Lysylphosphatidylglycerol synthase TM region | NA | Lysylphosphatidylglycerol synthase conveys resistance of *Staphylococcus aureus* against cationic antimicrobial peptides by modifying the cell membrane (Staubitz et al., 2004) |
| LppC | bacterial outer membrane antigens | NA | Bacterial surface proteins could be involved in recognition between symbionts and host |
| LRR | leucine-rich repeats | LRR: 6.08% (3688)  LRR_TYP: 5.05% (1263) | Present in obligate intracellular amoeba symbiont *Candidatus* Amoebophilus asiaticus genome, putative function in interactions with the host (Schmitz-Esser et al., 2010) |
| LTD | lamin tail domain | NA | Nuclear lamins are part of the nuclear lamina, probably also involved in DNA interaction (Stuurman et al., 1998) |
| LTXXQ | LTXXQ motif family protein | NA | Motif in CpxP, a member of the two-component signal transduction Cpx pathway, which reacts to cell envelope stresses and misfolded proteins (Zhou et al., 2011) |
| MIF | macrophage migration inhibitory factor | NA | Involved in innate immunity of vertebrates and probably of invertebrates, as well as adaptive immunity of vertebrates, also found in plants and a cyanobacterium with unclear function (Sparkes et al., 2017) |
| NARP1 | NMDA receptor-regulated protein 1 | NA | Eukaryotic domain with similarity to cell cycle regulating yeast acetyltransferase (Pfam) |
| NHL | NCL-1, HT2A and Lin-41, similarity to WD repeat | NA | Present in sponge microbiome metatranscriptomes, putative role in symbiont-host interactions (Díez-Vives et al., 2017) |
| Nipsnap | NIPSNAP | NA | Could be involved in vesicular trafficking (Lee et al., 2002)  Present in sponge microbiome metatranscriptomes, putative role in symbiont-host interactions (Díez-Vives et al., 2017) |
| PQQ, PQQ-like | PQQ enzyme repeat | 63.42% (6647) | Beta-propeller repeat in enzymes using pyrollo-quinoline quinone (PQQ) as prosthetic group (Pfam)  Present in sponge microbiome metatranscriptomes, putative role in symbiont-host interactions (Díez-Vives et al., 2017) |
| Sel1 | TPR subfamily | 61.04% (10391) | Present in sponge microbiome metatranscriptomes, putative role in symbiont-host interactions (Díez-Vives et al., 2017)  Present in obligate intracellular amoeba symbiont *Candidatus* Amoebophilus asiaticus genome, probable function in interactions with the host (Schmitz-Esser et al., 2010) |
| SNARE, IncA | SNARE domain, IncA protein | t_SNARE: 0.07% (4) | Intracellular bacteria can use SNARE-like proteins (including IncA) to block host SNARE-mediated membrane fusion and thereby endocytosis (Paumet et al., 2009) |
| TIR | Toll-interleukin-receptor | 1.04% (47) | Protein-protein interaction domain in innate immune system proteins of plants and animals, present in pathogenic and non-pathogenic bacteria, TIR domain-containing proteins of pathogens can interfere with host immune system (Ve et al., 2015) |
| TPR | tetratricopeptide repeats | 51.13% (33476) | Present in sponge microbiome metatranscriptomes, putative role in symbiont-host interactions (Díez-Vives et al., 2017)  Present in obligate intracellular amoeba symbiont *Candidatus* Amoebophilus asiaticus genome, probable function in interactions with the host (Schmitz-Esser et al., 2010)  TPR-containing bacterial proteins can have impact on phagocytosis of bacteria by amoebae (Reynolds and Thomas, 2016) |
| TSP | thrombospondins | TSP1: 0.15% (16)  TSPN: 0.18% (6) | Extracellular glycoproteins in animals, involved in cell attachment in vertebrates (Adams and Lawler, 2004) |
| TTL | tubulin-tyrosine ligase family | NA | Tubulin posttranslational modification (tyrosination), essential for cell development (Prota et al., 2013) |
| U-box | ubiquitin interacting | 0.44% (19) | Present in obligate intracellular amoeba symbiont *Candidatus* Amoebophilus asiaticus genome, could interact with host ubiquitination pathways (Schmitz-Esser et al., 2010)  Bacterial proteins might interact with ubiquitination pathways in the host (Rytkönen and Holden, 2007) |
| UB activating E1 | Domain of ubiquitin activating E1 family enzymes | UBA_E1_c: 0 | Bacterial proteins might interact with ubiquitination pathways in the host (Rytkönen and Holden, 2007) |
| UBA | ubiquitin associated domain | 0.61% (48) | Bacterial proteins might interact with ubiquitination pathways in the host (Rytkönen and Holden, 2007) |
| VIT | Vault protein Inter-alpha-Trypsin domain | 0 | Conserved domain in the inter-alpha-trypsin inhibitor heavy chain family, one of the precursor proteins of inter-alpha-trypsin inhibitors, which are protease inhibitors and involved in stabilization of the extracellular matrix (Himmelfarb et al., 2004) |
| VWA | Von Willebrand factor type A domain | 47.46% (20119) | Domain involved in eukaryotes in protein-protein interactions, multiprotein complexes, extracellular matrix, cell adhesion, various intracellular functions; prokaryotic proteins mostly not well characterized, characterized bacterial proteins with different functions; some prokaryotic VWA proteins possibly acquired via horizontal gene transfer (Whittaker and Hynes, 2002) |
| WD40, PD40 | WD40, WD40-like repeats | WD40: 5.54% (5471) | WD40 repeat proteins are highly abundant proteins in eukaryotes, involved in various functions; comparatively rare in prokaryotes, here putatively also involved in different functions, enriched in Cyanobacteria and Planctomycetes (Hu et al., 2017) |
