## Supplementary Table S7 for "Host-microbe interactions in the chemosynthetic *Riftia pachyptila* symbiosis"

Supp. Table S7: Transcriptome completeness for the four different assemblies based on the BUSCO eukaryote and metazoan datasets. C: complete BUSCOS, S: complete and single-copy BUSCOS, D: complete and duplicated BUSCOS, F: fragmented BUSCOS, M: missing BUSCOS. Assembly 4 was used for all further analyses.

|  | Completeness [%] | |
| --- | --- | --- |
| Assembly | *Eukaryota* | *Metazoa* |
| 1 | C: 99.0 [S: 32.3, D: 66.7], F: 1.0, M: 0.0 | C: 98.3 [S: 31.4, D: 66.9], F: 0.9, M: 0.8 |
| 2 | C: 99.0 [S: 33.0, D: 66.0], F: 0.7, M: 0.3 | C: 98.3 [S: 32.5, D: 65.8], F: 0.9, M: 0.8 |
| 3 | C: 99.0 [S: 36.0, D: 63.0], F: 1.0, M: 0.0 | C: 98.4 [S: 39.7, D: 58.7], F: 0.9, M: 0.7 |
| 4 | C: 99.3 [S: 33.0, D: 66.3], F: 0.7, M: 0.0 | C: 98.8 [S: 37.1, D: 61.7], F: 0.7, M: 0.5 |
