## Supplementary Table S8 for "Host-microbe interactions in the chemosynthetic *Riftia pachyptila* symbiosis"

| **Tool** | **Used for host/ symbiont (Sym) proteins** | **Prediction** | **Reference** |
| --- | --- | --- | --- |
| Pfam* | Host, Sym | functional annotation | (El-Gebali et al., 2019) |
| TIGRFAM* | Host, Sym | functional annotation | (Haft et al., 2001) |
| KEGG* | Host, Sym | functional annotation | (Kanehisa et al., 2017) |
| eggNOG* | Host, Sym | functional annotation | (Huerta-Cepas et al., 2016) |
| BlastKoala | Host, Sym | functional annotation | (Kanehisa et al., 2016) |
| Phobius | Host, Sym | signal peptides and transmembrane domains | (Käll et al., 2007) |
| SignalP | Host, Sym | signal peptides | (Nielsen, 2017) |
| TMHMM | Host, Sym | transmembrane helices | (Krogh et al., 2001) |
| CELLO | Host, Sym | subcellular localization | (Yu et al., 2006) |
| PSortb | Sym | subcellular localization | (Yu et al., 2010) |
| SecretomeP | Sym | non-classical secretion signals | (Bendtsen et al., 2005) |
| Lipo | Sym | lipoproteins | (Berven et al., 2006) |
| BOMP | Sym | β-barrel outer membrane proteins | (Berven et al., 2004) |
| TargetP | Host | subcellular localization | (Emanuelsson et al., 2000) |
| WoLF PSORT | Host | subcellular localization | (Horton et al., 2007) |
| DeepLoc | Host | subcellular localization | (Almagro Armenteros et al., 2017) |
| SLP-Local | Host | subcellular localization | (Matsuda et al., 2005) |

Supp. Table S8: Tools used to characterize *Riftia* host and symbiont proteins included in the combined *Riftia* host and symbiont database used in this study.

* Metaerg (https://sourceforge.net/projects/metaerg/) was used to query these databases
